# CCR4^+^low-density neutrophils define a novel immunosuppressive subset associated with breast cancer progression and early mortality

**DOI:** 10.64898/2026.09.22.753474

**Authors:** Bruna F. Correia, Daniela Grosa, Rute Salvador, Telma Martins, Marina Vitorino, Jorge M. Mendes, Andreia Lopes, Marta Parreira, Beatriz Martins, Guilherme Vilhais, Mariana Marques, Maria Gabriela Gasparinho, Sofia Critovão-Ferreira, Diana P. Saraiva, Nídia de Sousa, Sofia Braga, António Jacinto, M. Guadalupe Cabral

## Abstract

Neutrophils are increasingly recognized as key players in cancer progression, yet their functional heterogeneity limits their therapeutic exploitation. In breast cancer (BC), elevated circulating and tumor-associated neutrophils have been associated with poor prognosis, and low-density neutrophils (LDN), particularly, have been linked to immunosuppression. However, the specific neutrophil subsets driving these effects remain undefined. Here, we identify a previously unrecognized CCR4-expressing neutrophil subset, strongly enriched among LDN, whose frequency rises with disease stage and correlates with faster progression and shorter survival. Serial blood sampling further showed that CCR4^+^LDN levels fluctuate in line with clinical course, with increases indicating near-term risk in metastatic patients. Our findings also reveal that this subset has an immunosuppressive signature more pronounced than that of the remaining LDN, migrates preferentially toward tumor-derived chemokines, and dampens T cell-driven tumor killing even when PD-1 is blocked. Additionally, in a pilot cohort of Triple-Negative BC patients on anti-PD-1 therapy, early changes in this population were associated with treatment outcome. Together, these results reveal a distinct neutrophil population that actively contributes to immune escape and tumor progression, establishing CCR4^+^LDN as a minimally invasive real-time biomarker of disease course and treatment response, and as a promising target for neutrophil-directed therapies in BC.

**Graphical Abstract:** 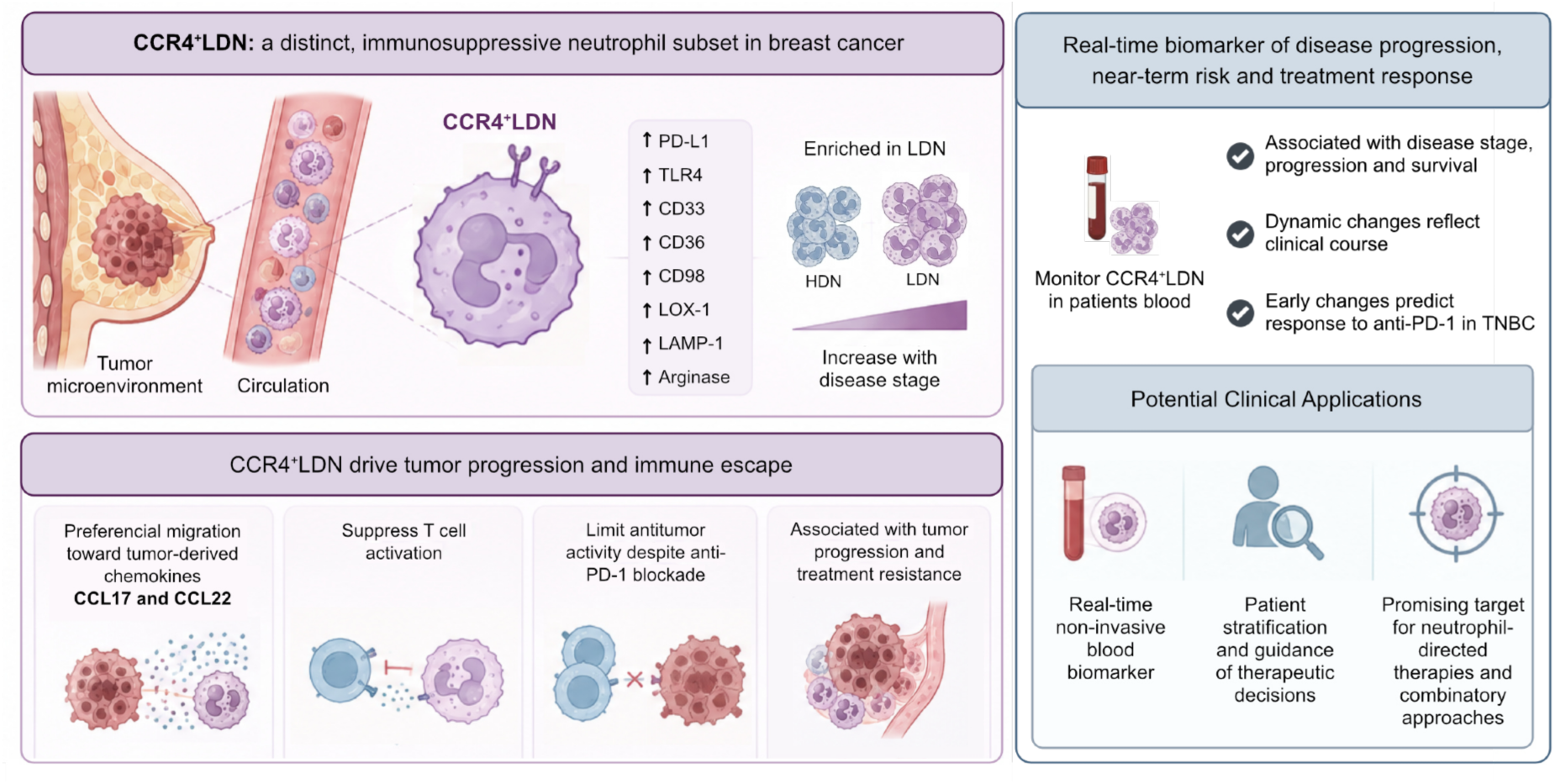

## Introduction

Breast cancer (BC) remains the most prevalent malignancy among women worldwide and a leading cause of cancer-related mortality (1), with five-year survival dropping to 25% once distant metastases develop (2). Among BC subtypes, triple-negative breast cancer (TNBC) represents the most aggressive form, characterized by the absence of hormone receptors and HER2 expression, limited targeted therapeutic options, and disproportionately high rates of recurrence, progression to metastatic disease, and mortality (3,4). Its immunogenic profile has positioned TNBC as the subtype most amenable to immune checkpoint blockade, and integrating this immunotherapy with chemotherapy has improved outcomes in this population, particularly in neoadjuvant regimens (5,6). However, although approximately 60% of patients with high-risk early-stage TNBC achieve a pathological complete response (pCR) after neoadjuvant pembrolizumab plus chemotherapy, patients who do not achieve pCR may have residual invasive disease and face a substantially increased risk of recurrence, distant metastasis, and death, despite the potential benefit of subsequent post-neoadjuvant treatment (7,8).

Growing evidence indicates that tumor progression and treatment resistance are determined not only by malignant cells but also critically shaped by the tumor microenvironment (TME), in which immune populations can either restrain or promote disease evolution (9,10,11,12).

Among the immune compartment, neutrophils have emerged as important yet incompletely understood regulators of cancer biology. Once considered short-lived and homogeneous, first responders, neutrophils are now recognized as highly plastic cells capable of adopting divergent phenotypes in response to tumor-derived signals (13,14,15). The early binary classification of tumor-associated neutrophils into antitumor (N1) and protumor (N2) states has been superseded by a more nuanced understanding of neutrophil heterogeneity, largely driven by single-cell technologies and patient-derived studies (16,17). In circulation, neutrophils can be stratified into high-density neutrophils (HDN) and low-density neutrophils (LDN), the latter being enriched in cancer and frequently associated with immunosuppressive and tumor-promoting functions (18,19,20), overlapping in some reports with the granulocytic fraction of myeloid-derived suppressor cells (G-MDSCs) (21). Elevated neutrophil counts and increased neutrophil-to-lymphocyte ratios consistently correlate with poor prognosis across multiple malignancies, including BC, underscoring their clinical relevance (22,23). Neutrophil recruitment to the TME is primarily mediated by chemokine signaling. This process is largely governed by CXCR1 and CXCR2 and their respective ligands — notably CXCL5, CXCL6, and CXCL8 — though CCR1 and CCR2 ligands have also been implicated (24). Whether LDN specifically use distinct or additional chemokine receptor repertoires to access the TME remains incompletely understood, highlighting a knowledge gap that may have important implications for their selective accumulation in cancer.

We have previously demonstrated that LDN accumulate in BC patients and associate with poor response to neoadjuvant chemotherapy and advanced disease (18,19). Functionally, LDN exhibit an enhanced capacity to suppress T cells’ activation through multiple mechanisms: direct cell-cell contact mediated by immune checkpoint ligands such as PD-L1, release of soluble immunosuppressive factors including reactive oxygen species and arginase, and even indirect immunosuppression via the recruitment of regulatory T cells (Tregs) through the release of CCL17 (18,19,25). Nevertheless, LDN represent a heterogeneous population, and the specific subsets contributing to disease progression remain poorly defined.

Here, we identify a previously unrecognized neutrophil subset expressing the chemokine receptor CCR4, markedly enriched within the LDN fraction in BC patients, particularly those with metastatic disease. While the CCR4–CCL17/CCL22 axis is a well-established mediator of Treg recruitment and peripheral tolerance (26), its role in neutrophil biology remained unexplored. We show that CCR4^+^LDN accumulate with advancing disease stage, associate with accelerated progression, and reduced survival.

Longitudinal analysis in the metastatic cohort further revealed that CCR4^+^LDN dynamics closely track clinical trajectories, with increasing frequencies providing greater information on near-term risk than baseline levels, supporting their potential utility for real-time disease monitoring. Additionally, in a pilot study of TNBC patients on anti-PD-1 therapy, early on-treatment changes in CCR4^+^LDN were associated with treatment response, suggesting that CCR4^+^LDN may serve as a dynamic, minimally invasive indicator of immunotherapy response and inform timely treatment adjustments.

Together, these findings define CCR4 as a marker of a functionally distinct neutrophil subset with direct clinical relevance and reveal an unexpected role for the CCR4–CCL17/CCL22 axis in neutrophil pathobiology. Our work provides a refined framework for understanding neutrophil heterogeneity in cancer and positions CCR4^+^LDN not only as a biomarker of disease progression and treatment response but also as a candidate therapeutic target to counteract immune evasion in BC.

## Results

### CCR4 expression identifies a distinct subset of circulating low-density neutrophils in breast cancer

To investigate the phenotypic heterogeneity of circulating neutrophils in BC, we performed flow cytometric analysis of LDN and HDN isolated by density gradient centrifugation from patients’ blood (Figure 1A). This analysis revealed a subset of circulating neutrophils expressing the chemokine receptor CCR4, detected in both LDN and HDN populations (Figure 1B and 1C). Although the frequency of CCR4-expressing neutrophils varied substantially among patients, CCR4 expression was consistently enriched within the LDN compartment, with significantly higher frequencies of CCR4^+^LDN (8.11% [IQR: 2.30 – 23.75%]) compared to HDN (0.65% [IQR: 0.14 – 2.00%]) across the cohort (Figure 1D). To our knowledge, CCR4 expression has not been previously reported in neutrophils. Collectively, these findings identify a previously unrecognized CCR4^+^ neutrophil population that is preferentially enriched within the circulating LDN compartment of patients with BC.

**Figure 1.**
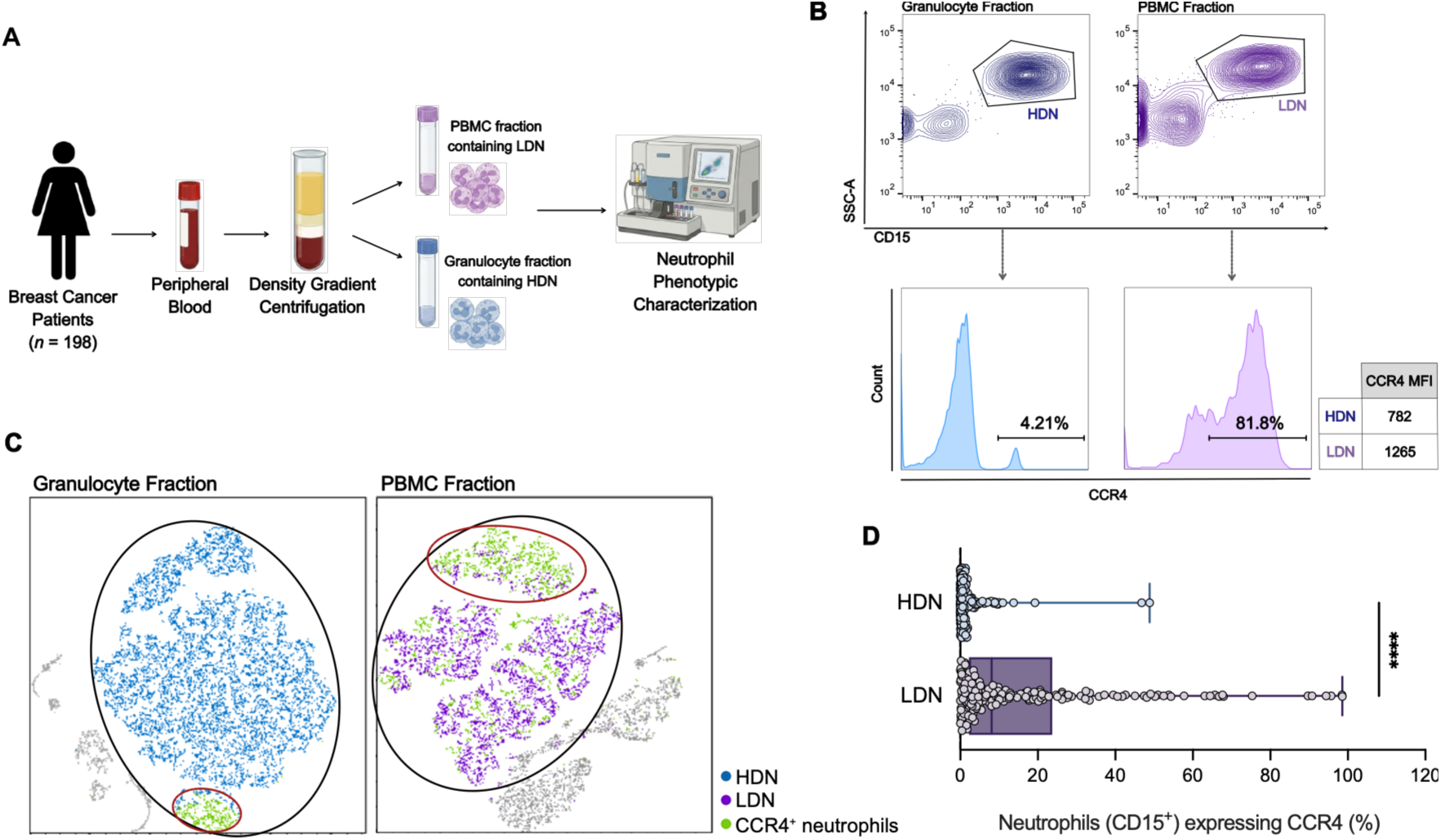
CCR4 expression identifies a novel subset of circulating neutrophils enriched within the low-density neutrophil compartment of patients with breast cancer. **(A)** Schematic overview of the experimental workflow. The PBMC (containing LDN) and granulocyte (containing HDN) fractions were obtained by density gradient centrifugation and analyzed by flow cytometry. **(B)** Representative gating strategy identifying CD15^+^ neutrophils in each fraction and representative histograms illustrating CCR4 expression in HDN and LDN. **(C)** Representative t-SNE visualization of independently acquired PBMC and granulocyte fractions, highlighting the expansion and distribution of CCR4⁺ neutrophils within each compartment. **(D)** Frequency of CCR4^+^ neutrophils among the HDN (blue, *n* = 174) and LDN (purple, *n* = 181) subpopulations from patients with BC, quantified by flow cytometry. \*\*\*\**p* < 0.0001, calculated with a Mann-Whitney test.

### CCR4^+^LDN accumulate during breast cancer progression and predict early mortality in metastatic disease

Having identified CCR4 as a marker of a previously unrecognized LDN subset, we next investigated its clinical relevance in BC. CCR4^+^LDN frequencies were quantified by flow cytometry in peripheral blood samples from healthy donors and patients with non-metastatic or metastatic BC. Consistent with our previous observations regarding total LDN (19), CCR4^+^LDN were virtually absent in healthy donors but significantly increased in patients with non-metastatic and metastatic BC (Figure 2A). Notably, metastatic patients exhibited significantly higher frequencies of CCR4^+^LDN (1.46% [IQR: 0.37 – 3.68%]) compared with non-metastatic patients (0.74% [IQR: 0.18 – 1.97%]), indicating a progressive expansion of this subset with disease advancement (Figure 2A). Importantly, CCR4^+^LDN frequencies were not specifically associated with a BC subtype, nor with the number or anatomical site of metastasis (bone, visceral, or nervous system (CNS)), with comparable interpatient heterogeneity observed across all four subtypes and metastatic sites (Supplemental Figure 1). A multivariable model further showed that, among metastatic patients, CCR4^+^LDN frequency did not increase with molecular subtype, number of metastatic sites, visceral involvement, serum tumor marker levels, or disease setting (relapse *vs* de novo) (Supplemental Table 1). Altogether, these observations indicate that CCR4^+^LDN is a broad marker of advanced disease rather than one that is further graded by metastatic burden.

**Figure 2.**
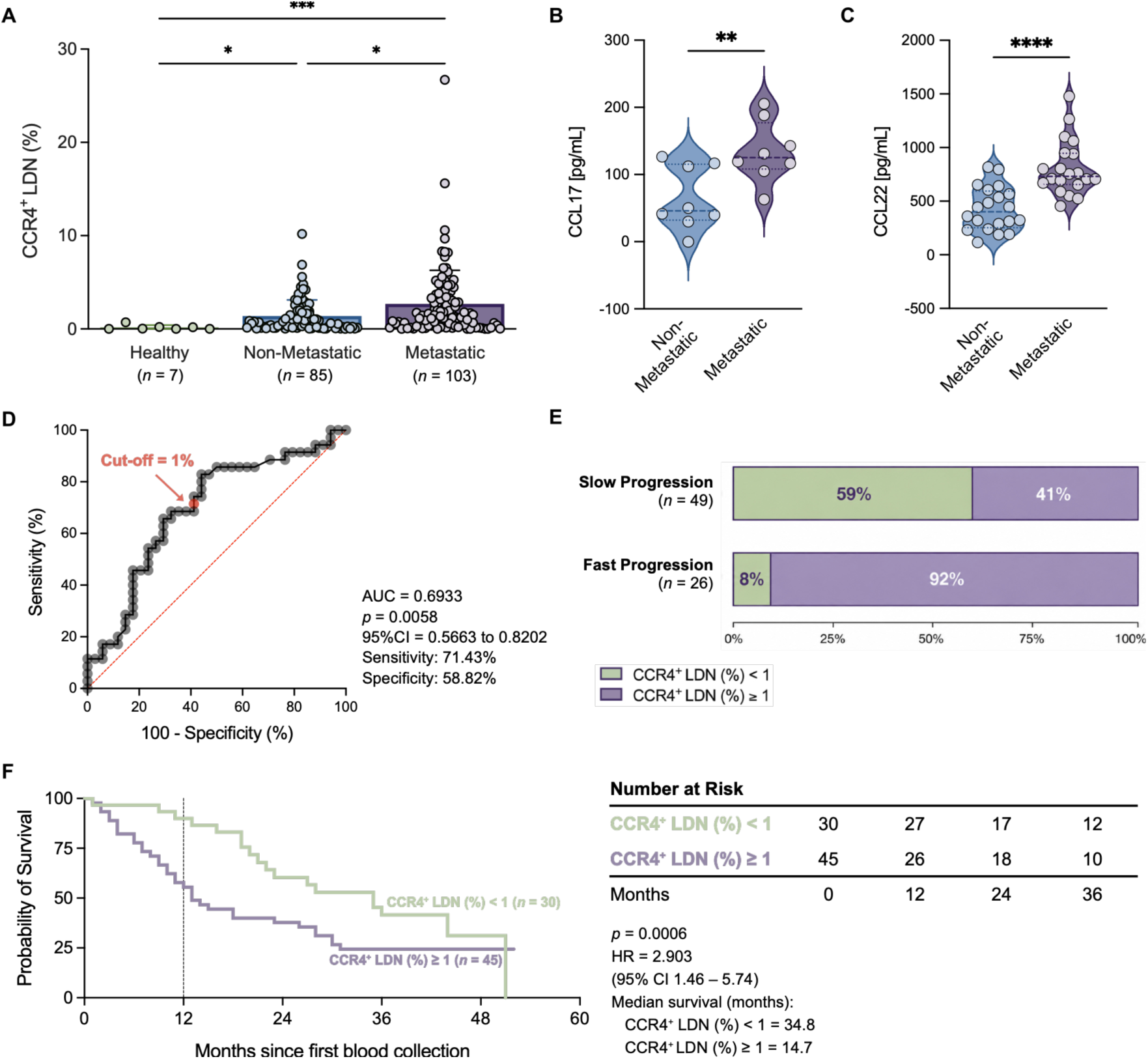
CCR4^+^LDN expand during breast cancer progression and are associated with poor clinical outcomes. **(A)** Frequency of circulating CCR4^+^LDN in healthy donors (green, *n =* 7), non-metastatic BC (blue, *n* = 85), and metastatic BC patients (purple, *n* = 103). **(B and C)** Plasma concentrations of the CCR4 ligands CCL17 and CCL22 in non-metastatic and metastatic BC patients, measured by ELISA. **(D)** Receiver operating characteristic (ROC) curve analysis used to determine the optimal cut-o^ for CCR4^+^LDN frequency within the PBMC fraction. **(E)** Distribution of metastatic BC patients with slow progression (*n* = 49) and rapid progression (*n* = 26), according to CCR4^+^LDN frequency at the time of the first blood collection for our study. Rapid progression was defined as disease progression or death within 12 months after sample collection. **(F)** Kaplan-Meier survival analysis of metastatic BC patients stratified according to CCR4^+^LDN frequency in the PBMC fraction. Green line, CCR4^+^LDN <1% (*n* = 30); purple line, CCR4⁺LDN ≥1% (*n* = 45). \*\*\*\**p* < 0.0001, \*\*\**p* < 0.001, \*\**p* < 0.01, calculated with a Kruskal-Wallis test with Dunn’s multiple comparisons test (A), a Welch Test (B and C), or a Gehan-Breslow-Wilcoxon test (D).

Next, we assessed the circulating levels of CCR4 ligands, CCL17 and CCL22, in different stages of BC. Plasma concentrations of both chemokines were significantly higher in metastatic BC patients (CCL17: 134.00 ± 45.47 pg/mL; CCL22: 809.20 ± 257.30 pg/mL) than in those with non-metastatic disease (CCL17: 64.61 ± 47.13 pg/mL; CCL22: 428.60 ± 204.20 pg/mL) (Figure 2B and 2C). This pattern mirrored the increased frequency of CCR4^+^LDN observed in metastatic patients, supporting an association between the CCR4-CCL17/22 axis and the expansion of this neutrophil subset during disease progression.

We also evaluated the prognostic significance of circulating CCR4^+^LDN in metastatic BC. Receiver operating characteristic (ROC) curve analysis identified an optimal cutoff of 1% CCR4^+^LDN (within the PBMC fraction), maximizing sensitivity while maintaining an acceptable specificity to minimize false negatives, which was subsequently used to stratify patients for disease progression and survival analyses (Figure 2D). Patients with CCR4^+^LDN frequencies ≥1% experienced significantly more rapid disease progression, defined as disease worsening or death within one year after sample collection, than patients with lower CCR4^+^LDN frequencies (Figure 2E). Kaplan-Meier analysis further demonstrated that BC patients with CCR4^+^LDN frequencies ≥1% had shorter overall survival than those with lower frequencies (median 14.7 *vs* 34.8 months, respectively) (Figure 2F). Notably, the prognostic impact of CCR4^+^LDN was time-dependent, with a significant violation of the proportional hazards assumption (Schoenfeld residuals test, *p* = 0.0007) (Supplemental Figure 2). Time-varying Cox regression showed that CCR4^+^LDN frequencies ≥1% were strongly associated with mortality during the first 12 months after sample collection (HR = 5.08, 95% CI 1.5–17.2, *p* = 0.009), but not thereafter (HR = 0.87, 95% CI 0.42–1.81, *p* = 0.705) (Supplemental Table 2, Figure 2F). This early association remained evident after stratification by metastatic site, with significantly reduced survival among patients with bone and visceral metastases and a similar trend for CNS metastases (Supplemental Figure 2).

Collectively, these observations indicate that CCR4^+^LDN represent a clinically relevant neutrophil subset that expands during BC progression and is associated with poor clinical outcome in metastatic disease. Importantly, elevated CCR4^+^LDN frequencies identify patients at particularly high risk of early disease progression and death, highlighting their potential as a prognostic biomarker in BC.

### Longitudinal CCR4^+^LDN dynamics track metastatic breast cancer course and predict real-time mortality risk

Having established the prognostic significance of CCR4^+^LDN, we next investigated whether longitudinal changes in this subset reflected disease evolution during follow-up. CCR4^+^LDN frequencies were serially quantified in 46 metastatic BC patients with available longitudinal samples, and their dynamics were compared with treatment response and disease progression (Figure 3A). Patients whose CCR4^+^LDN frequencies remained below the 1% threshold generally experienced sustained disease control, whereas persistently elevated or increasing CCR4^+^LDN levels frequently coincided with disease progression, treatment failure, or death (Figure 3A).

**Figure 3.**
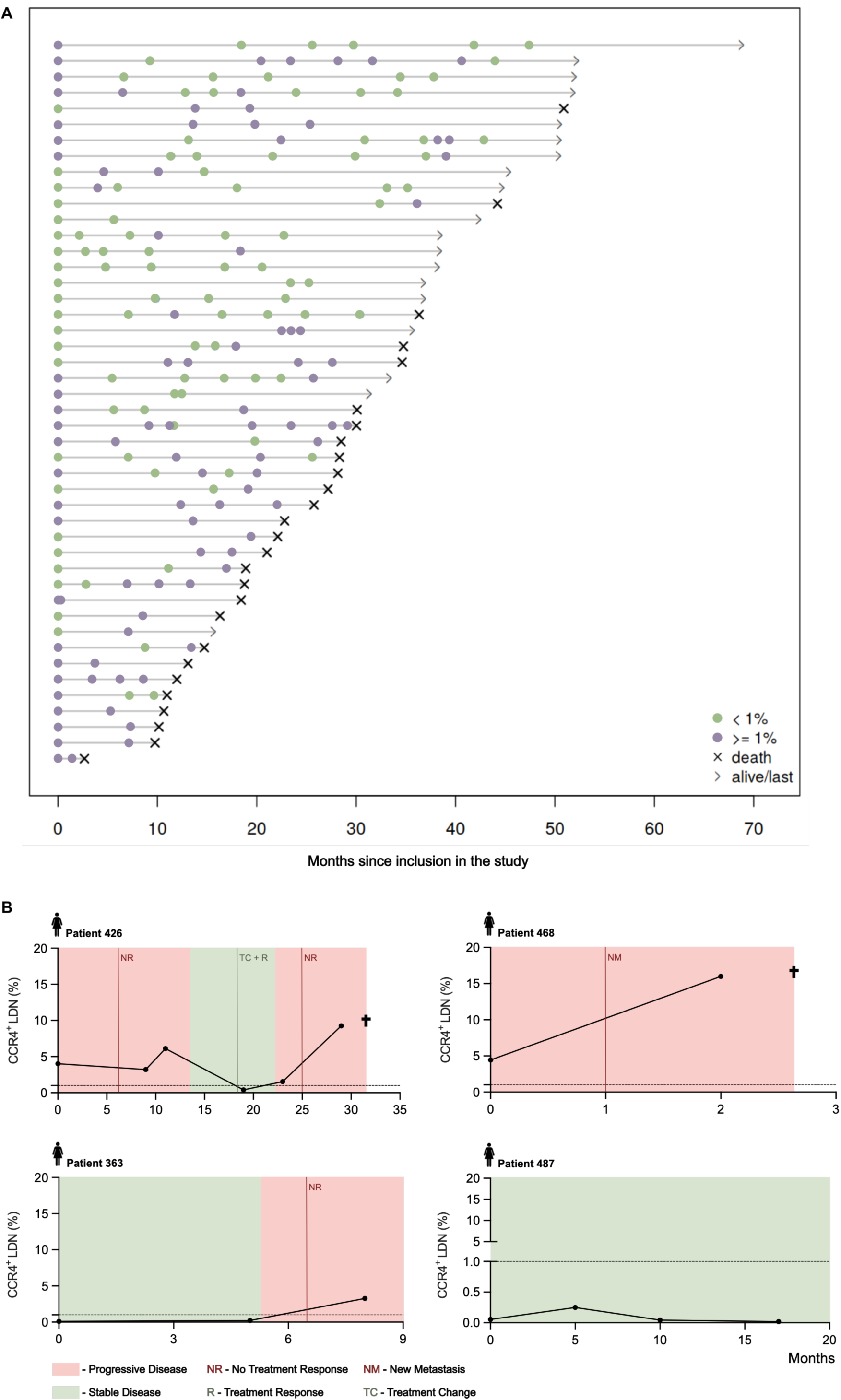
Fluctuations in CCR4^+^ low-density neutrophils are associated with changes in the clinical status of metastatic breast cancer. **(A)** Swimmer plot illustrating the percentage of CCR4^+^ low-density neutrophils (CCR4^+^LDN), assessed within the peripheral blood mononuclear cell (PBMC) layer, of 46 metastatic breast cancer (mBC) patients and respective relevant clinical events over time. Each bar represents a patient, with its length corresponding to the study enrollment period (in months). The coloured dots reflect the CCR4^+^LDN percentages in the blood (green -CCR4^+^LDN < 1%; purple -CCR4^+^LDN ≥ 1%). **(B)** Illustrative cases displaying longitudinal fluctuations in CCR4^+^LDN percentages over time in relation to disease status and clinical outcomes for four specific mBC patients.

Representative patient trajectories further illustrated that increases in circulating CCR4^+^LDN often preceded or accompanied the appearance of new metastatic lesions and clinical deterioration, whereas persistently low frequencies were associated with favourable treatment responses and prolonged disease stability (Figure 3B).

To determine whether the current, serially updated CCR4^+^LDN value carried greater prognostic information than the value measured at enrolment, we modelled CCR4^+^LDN as a time-varying covariate in the Cox regression. The current value was significantly associated with mortality risk (HR ≈ 1.6 per log-unit increase; ≥1% threshold associated with a ≈ 4-fold increase in risk), whereas the baseline (enrolment) value alone was a substantially weaker predictor of long-term outcome (Supplemental Table 3). This finding is consistent with the time-restricted prognostic effect observed in the earlier survival analysis (Figure 2F), and indicates that the most recently measured CCR4^+^LDN value, rather than the baseline value, best reflects a patient’s current risk.

Altogether, these results establish CCR4^+^LDN as a dynamic biomarker whose serially updated value tracks a patient’s risk of death in real time, more informatively than a single baseline measurement. This behaviour positions CCR4^+^LDN not simply as a static prognostic marker but as a longitudinal monitoring biomarker.

### CCR4^+^LDN display a distinct immunosuppressive and protumor phenotype

Given the clinical relevance of CCR4^+^LDN, we next sought to define the phenotypic characteristics distinguishing this subset from the remaining circulating LDN population. CCR4^+^ and CCR4^-^LDN were therefore compared using an expanded flow cytometry panel of markers associated with neutrophil maturation, activation, metabolism, and immunosuppressive function (Figure 4A, 4B and 4D). Compared with CCR4^-^LDN, the CCR4^+^LDN subset exhibited significantly higher frequencies of cells expressing CD33, PD-L1, TLR4, CD36, LOX-1, CD98, and LAMP-1 (Figure 4A and 4B). These findings establish CCR4^+^LDN as a more immature, metabolically active, and immunosuppressive neutrophil subset than their CCR4^-^counterparts.

**Figure 4.**
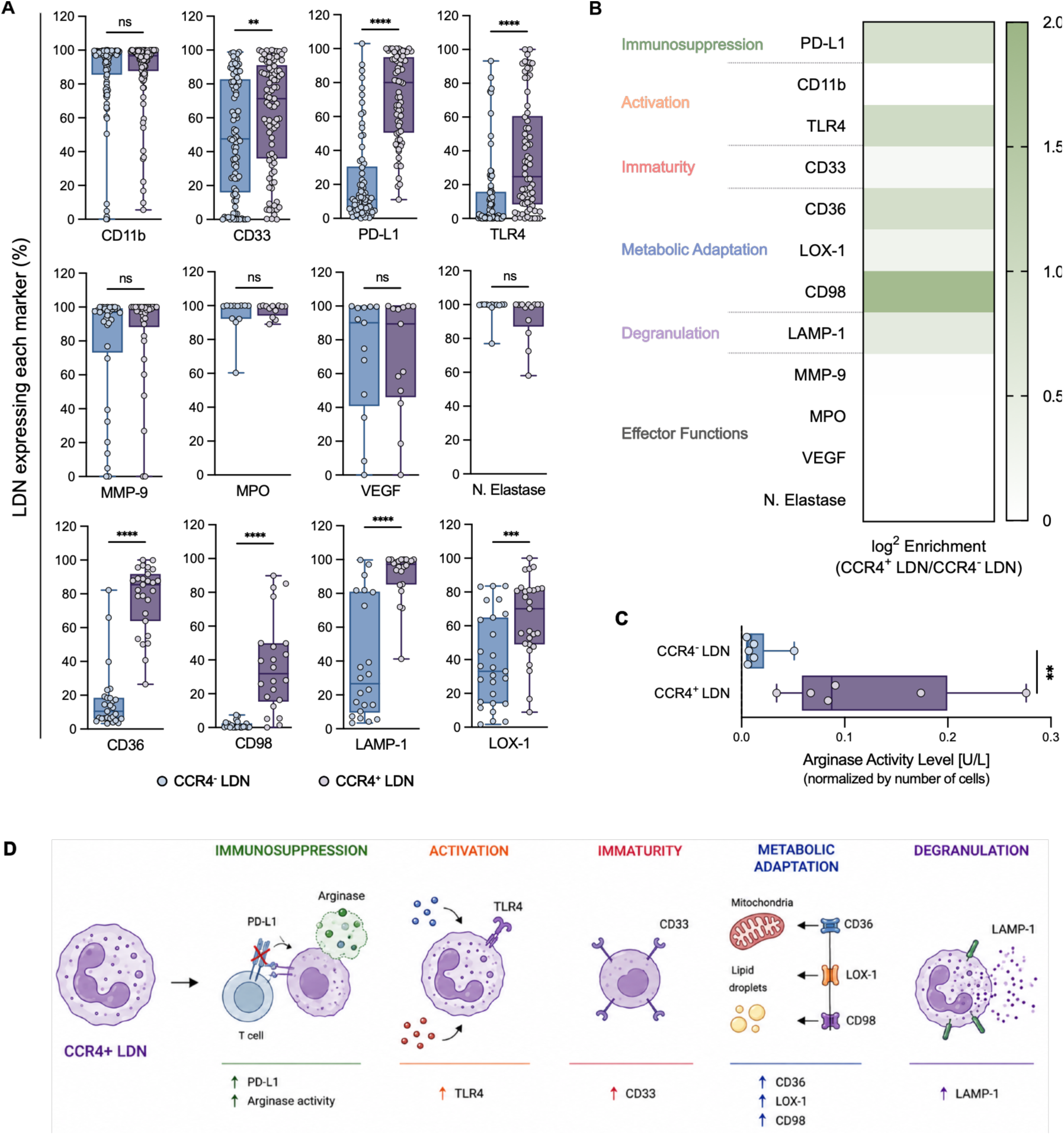
CCR4^+^LDN exhibit an immunosuppressive, activated, immature, and metabolically adapted phenotype. **(A)** Percentage of low-density neutrophils not expressing CCR4 (CCR4^-^LDN, blue bars) and LDN expressing CCR4 (CCR4^+^LDN, purple bars) (both assessed within the PBMC layer) expressing CD11b (*n* = 95), CD33 (*n* = 93), PD-L1 (*n* = 70), TLR4 (*n* = 70), MMP-9 (*n* = 34), MPO (*n* = 13), VEGF (*n* = 13), neutrophil elastase (*n* = 13), CD36 (*n* = 27), CD98 (*n* = 22), LAMP-1 (*n* = 22) and LOX-1 (*n* = 27) assessed by flow cytometry. **(B)** Summary heatmap showing the relative enrichment (log^2^ fold change) of phenotypic markers in CCR4^+^LDN compared with CCR4^-^LDN, highlighting pathways associated with immunosuppression, activation, immaturity, metabolic adaptation, degranulation, and e^ector functions. **(C)** Arginase activity levels measured in CCR4^+^LDN and CCR4^-^LDN (*n* = 6). **(D)** Schematic summary of the phenotypic characteristics enriched in CCR4^+^LDN. Statistical significance was determined using the Mann–Whitney test (A and C). \*\**p* < 0.01, *** *p* < 0.001, **** *p* < 0.0001.

Increased CD33 expression supports a less differentiated phenotype, whereas enrichment for PD-L1 suggests an enhanced capacity to suppress T cell responses. Elevated expression of TLR4, CD36, LOX-1, CD98, and LAMP-1 further indicates increased responsiveness to inflammatory and tumor-derived signals, along with metabolic adaptation and cellular activation. In contrast, no significant differences were observed in the frequencies of cells expressing CD11b, MMP-9, MPO, VEGF, or neutrophil elastase between CCR4^+^ and CCR4^-^LDN (Figure 4A and 4B).

Based on our previous observations that arginase activity is one of the mechanisms underlying LDN-mediated immunosuppression (19), we assessed whether this pathway was particularly relevant in CCR4^+^LDN. CCR4^+^LDN exhibited significantly higher arginase activity (0.0875 U/L [IQR: 0.0588 – 0.1995]) than CCR4^-^LDN (0.0095 U/L [IQR: 0.0058 – 0.0218]) (Figure 4C), further supporting an enhanced immunosuppressive capacity for this subset.

As such, together these findings demonstrate that CCR4 identifies a phenotypically and functionally distinct LDN subset enriched for features associated with immunosuppression and metabolic adaptation.

### TGF-β signaling is linked with CCR4 expression in neutrophils via a FOXP3-associated mechanism

We next sought to investigate the mechanisms regulating CCR4 expression in neutrophils. Given that TGF-β induces CCR4 expression in Tregs through a FOXP3-dependent pathway (26), and that this cytokine is also a key driver of neutrophil polarization towards a protumor phenotype (27), we assessed whether a similar mechanism could operate in neutrophils. We first confirmed that plasma TGF-β concentration levels increased with disease progression, being significantly higher in metastatic (21022 pg/mL, [IQR 8535 – 30110 pg/mL]) than in non-metastatic (6197 pg/mL, [IQR 3772 – 10047 pg/mL]) BC patients (Figure 5A). Moreover, circulating TGF-β levels positively correlated with the frequency of CCR4^+^LDN (Figure 5B), consistent with a potential role for TGF-β in the expansion of this neutrophil subset.

**Figure 5.**
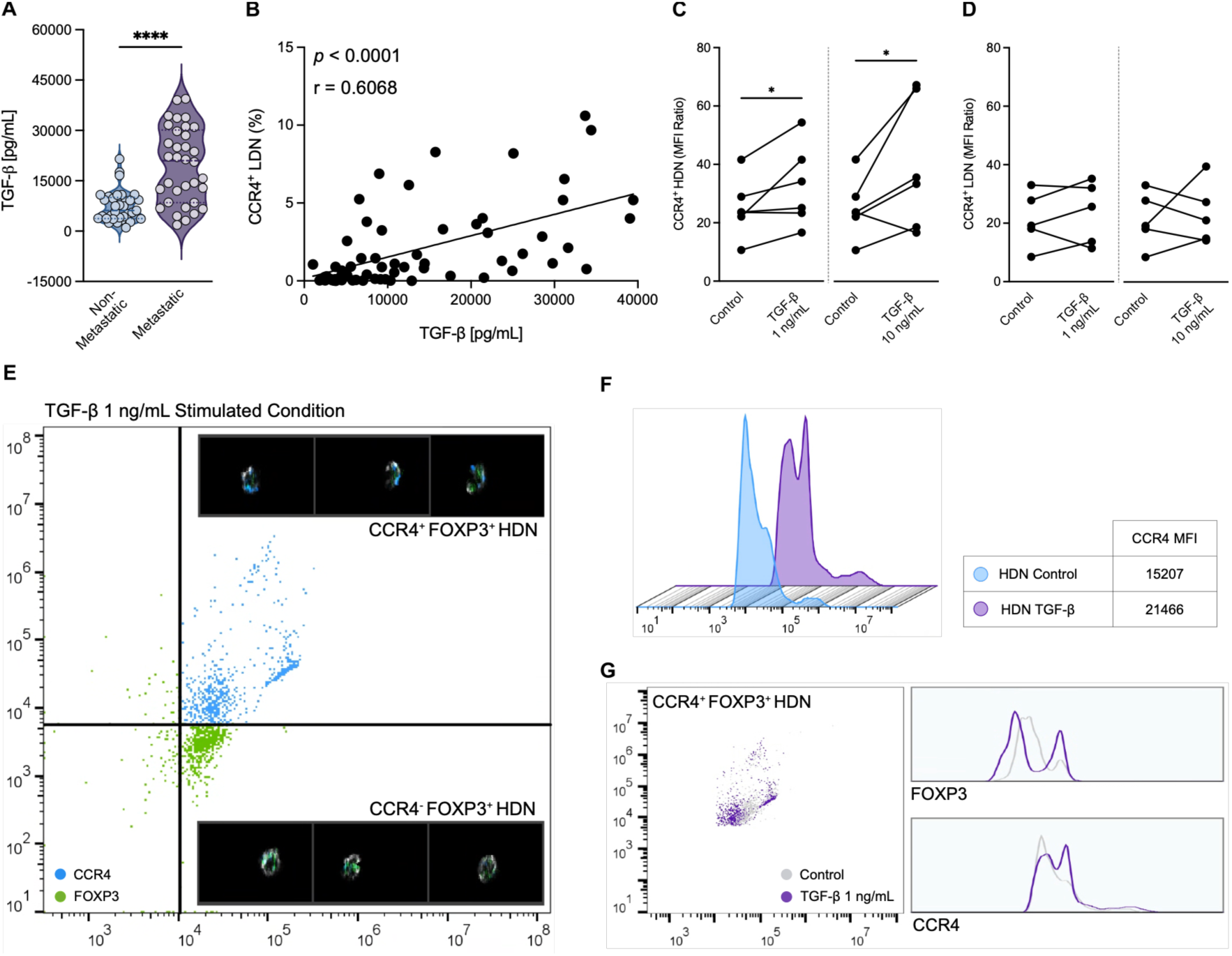
TGF-β signaling is associated with CCR4 expression in neutrophils through a FOXP3-associated mechanism. **(A)** Plasma TGF-β levels in non-metastatic (blue, *n* = 33) and metastatic (purple, *n* = 34) BC patients. **(B)** Correlation between plasma TGF-β levels and the frequency of circulating CCR4^+^LDN in BC patients (*n* = 62). **(C)** CCR4 expression in HDN from BC patients following stimulation with TGF-β (1 or 10 ng/mL) or control, expressed as the CCR4 MFI ratio (*n* = 5). **(D)** CCR4 expression in LDN from BC patients following stimulation with TGF-β (1 or 10 ng/mL) or control, expressed as the CCR4 MFI ratio (*n* = 5). **(E)** Representative imaging flow cytometry analysis showing co-expression of CCR4 and FOXP3 in individual HDN following TGF-β stimulation, with representative images of CCR4^+^FOXP3^+^ and CCR4^-^FOXP3^+^ neutrophils. **(F)** Representative CCR4 expression profiles in HDN before and after TGF-β stimulation, with corresponding CCR4 MFI values. **(G)** Representative flow cytometry analysis and fluorescence intensity profiles showing FOXP3 and CCR4 expression in HDN following TGF-β stimulation. Statistical significance was determined using a Mann–Whitney test (A), Spearman’s correlation (B), or a paired t-test (C, D and E). \**p* < 0.05, **** *p* < 0.0001.

We therefore examined whether TGF-β directly regulates CCR4 expression in neutrophils. TGF-β treatment significantly increased CCR4 expression in HDN (“normal” neutrophils) (Figure 5C), whereas no significant changes were observed in LDN (Figure 5D). These findings suggest that TGF-β contributes to CCR4 expression during neutrophil polarization rather than driving the expansion of already-committed protumor cells.

Considering that FOXP3 mediates TGF-β-driven CCR4 expression in Tregs, we next investigated whether neutrophils also express this transcription factor. Imaging flow cytometry further confirmed the co-expression of FOXP3 and CCR4 within individual neutrophils (Figure 5E). Notably, TGF-β stimulation increased the number of cells expressing both FOXP3 and CCR4, as well as the expression of CCR4 in HDN (Figure 5F and 5G), supporting a potential mechanistic link between TGF-β signaling, FOXP3 activation, and CCR4 induction.

### CCR4^+^LDN display enhanced migratory capacity toward breast cancer–associated chemotactic cues

Considering the role of CCR4 in Treg recruitment to the tumor, we investigated whether it also contributes to neutrophil migratory capacity. LDN exhibited significantly greater migration than HDN in response to both CCL17 and CCL22 (Supplemental Figure 4). Moreover, within the migrated LDN population, CCR4^+^ neutrophils were significantly enriched compared with CCR4^-^cells (Figure 6A and 6B). To determine whether this migratory advantage was mediated by CCR4, LDN were pre-incubated with a CCR4-blocking antibody prior to migration. CCR4 blockade significantly reduced LDN migration toward both CCL17 and CCL22 (Figure 6A and 6C), demonstrating that this response is CCR4-dependent. We next investigated whether BC cell lines similarly promote preferential recruitment of CCR4^+^LDN. Using MDA-MB-231 cells as the chemoattractant, CCR4^+^LDN displayed significantly greater migration than CCR4^-^LDN (Figure 6D and 6E). Importantly, CCR4 blockade markedly reduced neutrophil migration toward MDA-MB-231 cells (Figure 6D and 6F), indicating that CCR4 also contributes to tumor-promoted recruitment of LDN.

**Figure 6.**
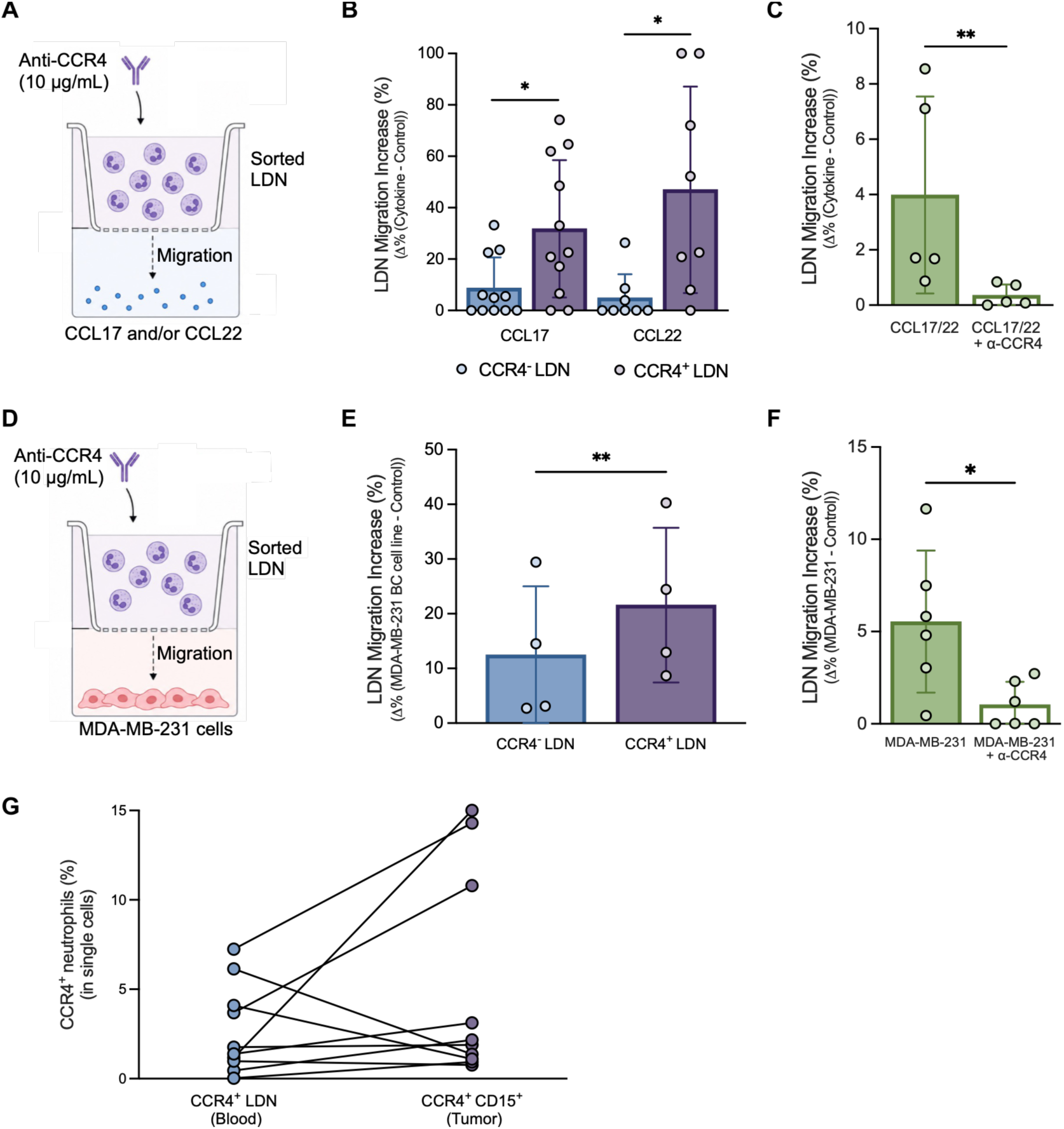
CCR4 mediates LDN migration toward BC-associated chemotactic cues, and CCR4^+^ neutrophils are detected within breast tumors. **(A)** Schematic overview of the transwell migration assays with CCL17 and CCL22 recombinant proteins. **(B)** Percentage increase in migration of CCR4^-^LDN (blue) and CCR4^+^LDN (purple) toward CCL17 (*n* = 11) and CCL22 (*n* = 8) relative to the control. **(C)** Percentage increase in migration of non-treated LDN (green, *n* = 5) and LDN treated with anti-CCR4 blocking (α-CCR4, white, *n* = 5) toward CCL17 and CCL22 relative to the control. **(D)** Schematic overview of the transwell migration assays with the MDA-MB-231 BC cell line. **(E)** Percentage increase in migration of CCR4^-^LDN (blue, *n* = 4) and CCR4^+^LDN (purple, *n* = 4) toward MDA-MB-231 cells relative to the control. **(F)** Percentage increase in migration of non-treated LDN (green, *n* = 6) and LDN (white, *n* = 6) treated with anti-CCR4 blocking (α-CCR4) toward MDA-MB-231 cells relative to the control. **(G)** Frequencies of CCR4^+^LDN (blue) and CCR4^+^ neutrophils (purple) assessed in matched peripheral blood (within the PBMC layer) and tumor samples, respectively (*n* = 10). Statistical significance was determined using a Wilcoxon matched-pairs signed-rank test (B and C), or a paired t-test (E and F). \**p* < 0.05, ** *p* < 0.01.

Supporting these data, CCR4^+^ neutrophils were indeed detected in dissociated breast tumor specimens (Figure 6G). In matched peripheral blood and tumor samples, patients with detectable circulating CCR4^+^LDN also showed CCR4^+^ neutrophils infiltrating their tumors (Figure 6G). Although evaluated in a limited number of patients, these findings indicate that CCR4^+^ neutrophils are present within the breast tumor microenvironment and are consistent with recruitment of this subset from the circulation.

Together, these results identify CCR4 as a functional mediator of LDN trafficking toward breast cancer-associated chemotactic cues and support a role for the CCR4-CCL17/CCL22 axis in the preferential recruitment of this immunosuppressive neutrophil subset to the tumor microenvironment.

### CCR4^+^LDN suppress T cell activation and antitumor activity despite PD-1 blockade

To determine whether the phenotypic differences observed in CCR4^+^LDN also translated into functional immunosuppressive activity, we evaluated their ability to modulate T cell response. Purified CCR4^+^ or CCR4^-^LDN were co-cultured with autologous CD15-depleted PBMCs, followed by stimulation with PMA and ionomycin. Exposure to CCR4^+^LDN significantly reduced the expression of T cell activation markers compared with CCR4^-^LDN (Figure 7A), indicating that this subset exerts a stronger suppressive effect on T cell activation. These findings are consistent with the increased PD-L1 expression and arginase activity observed in CCR4^+^LDN and further support their enhanced immunosuppressive phenotype.

**Figure 7.**
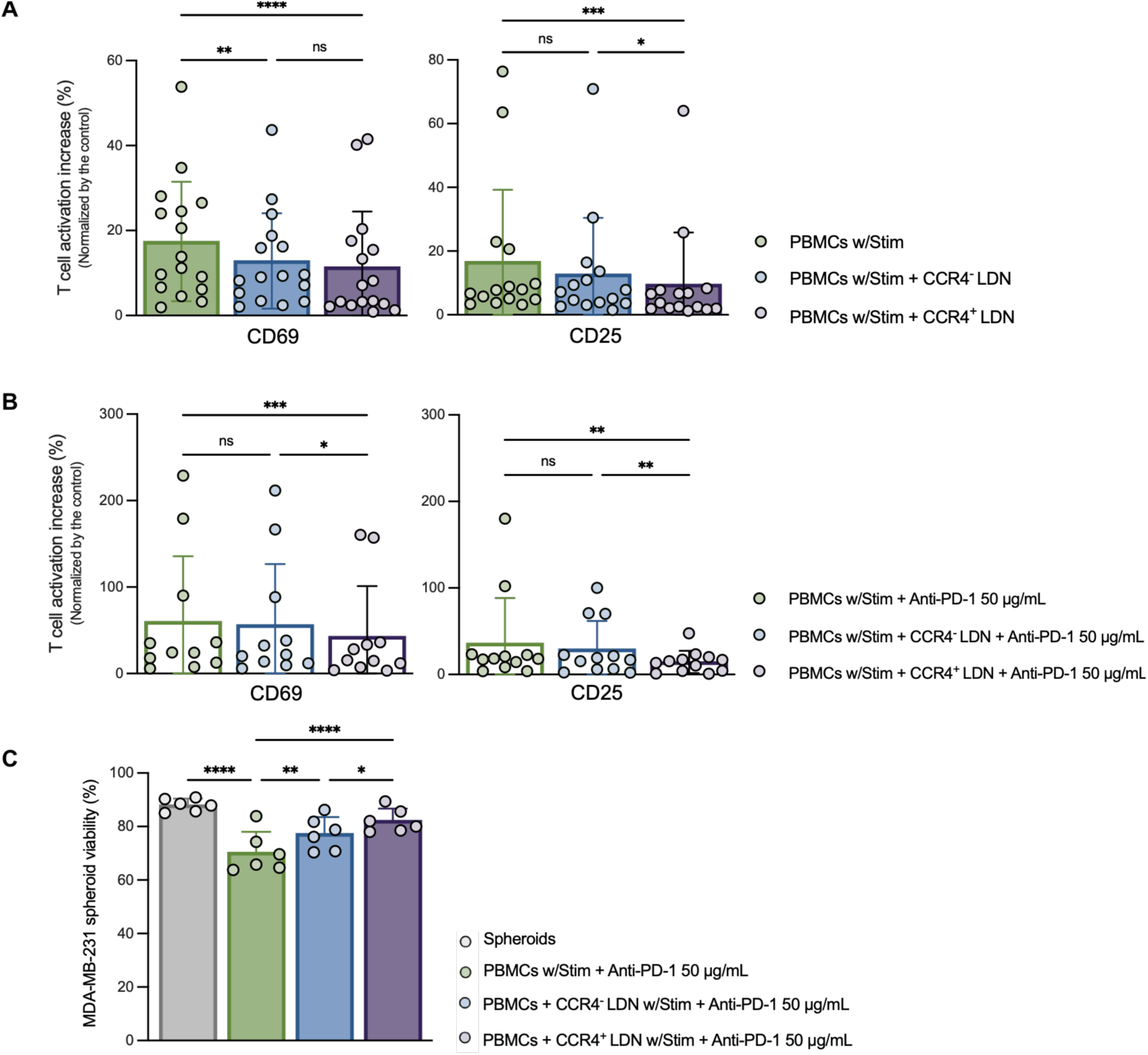
CCR4^+^ LDN impair T cell activation and T cell-mediated antitumor activity despite PD-1 blockade. **(A)** Percentage increase in CD69 and CD25 expression in stimulated CD3^+^ T cells cultured alone (green) or following co-culture with CCR4^-^LDN (blue) or CCR4^+^LDN (purple), normalized to the control condition (*n* = 16). **(B)** Percentage increase in CD69 and CD25 expression in the presence of anti-PD-1 blockade in stimulated CD3^+^ T cells cultured alone (green) or following co-culture with CCR4^-^LDN (blue) or CCR4^+^LDN (purple), normalized to the control condition (*n* = 11). **(C)** Viability of MDA-MB-231 tumor spheroids cultured alone (grey) or co-cultured with stimulated T cells (green) or with T cells previously stimulated in the presence of CCR4^-^LDN (blue) or CCR4^+^LDN (purple). Anti-PD-1 was added during tumor spheroid co-culture (*n* = 6). Statistical significance was determined using a Friedman test followed by Dunn’s multiple-comparisons test (A and B), or a repeated-measures one-way ANOVA followed by Holm–Šidák multiple-comparison test (C). \**p* < 0.05, ** *p* < 0.01, *** *p* < 0.001, **** *p* < 0.0001.

We next investigated whether PD-1 blockade could overcome this suppressive effect. Although treatment with a clinically used anti-PD-1 agent improved T cell activation in addition to stimulation, this improvement failed to fully overcome the suppression induced by CCR4^+^LDN, as T cells previously exposed to CCR4^+^LDN remained significantly less activated than those exposed to CCR4^-^LDN (Figure 7B).

To determine whether this impaired activation translated into reduced antitumor function, activated T cells pre-conditioned by CCR4^+^ or CCR4^-^LDN were co-cultured with MDA-MB-231 tumor spheroids in the presence of anti-PD-1. Again, T cells exposed to CCR4^+^LDN displayed a reduced capacity to eliminate tumor cells, resulting in significantly higher spheroid viability compared with T cells conditioned by CCR4^-^LDN (Figure 7C).

Together, these findings demonstrate that CCR4^+^LDN suppress T cell activation and compromise T cell-mediated antitumor activity despite PD-1 blockade, supporting a potential role for this neutrophil subset in limiting the efficacy of PD-1-targeted immunotherapy in BC.

### Circulating CCR4^+^LDN dynamics associate with clinical response to anti-PD-1 therapy in triple-negative breast cancer

Considering the enhanced immunosuppressive activity of CCR4^+^LDN and its association with worse prognosis in BC, we next investigated the clinical relevance of this neutrophil subset in patients receiving anti-PD-1 therapy. To this end, we analyzed a prospective cohort of patients with TNBC treated with pembrolizumab combined with chemotherapy in the neoadjuvant setting and longitudinally monitored circulating CCR4^+^LDN frequencies throughout treatment (Figure 8A). Although baseline CCR4^+^LDN frequencies showed limited discrimination between responders and non-responders, evaluation at the first on-treatment follow-up revealed a separation between the two groups, with responders displaying significantly lower CCR4^+^LDN frequencies (0.46% [IQR: 0.31 – 0.79%]) than non-responders (1.47% [IQR: 1.04 – 3.33]) (Figure 8B and 8C), consistent with the previous *in vitro* results.

**Figure 8.**
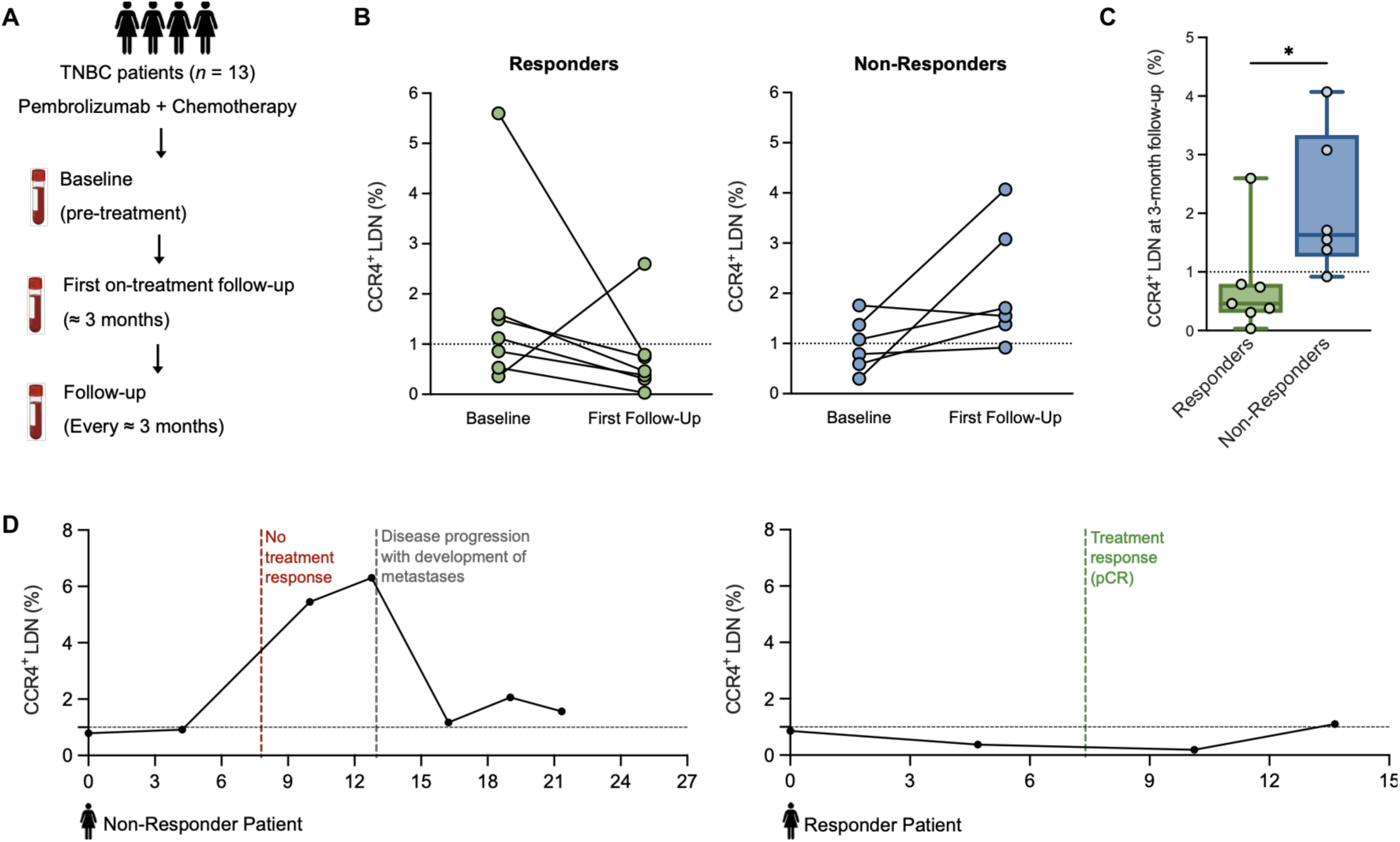
Longitudinal circulating CCR4^+^LDN dynamics are associated with clinical response to anti-PD-1 therapy in triple-negative breast cancer. **(A)** Schematic representation of the prospective longitudinal monitoring of circulating CCR4^+^LDN in patients with TNBC treated with pembrolizumab in combination with chemotherapy in the neoadjuvant setting. **(B)** Frequency of circulating CCR4^+^LDN at baseline and at first follow-up in TNBC patients subsequently classified as responders (green, *n* = 7) or non-responders (blue, *n* = 6). **(C)** Frequency of circulating CCR4^+^LDN at the first early on-treatment follow-up in responders (green, *n* = 7) and non-responders (blue, *n* = 6). **(D)** Longitudinal changes in CCR4^+^LDN frequencies in a representative responder and non-responder patient throughout treatment, shown in relation to their clinical trajectories. Statistical significance was determined using a Mann-Whitney test (C). *p < 0.05.

This indicates that early changes in circulating CCR4^+^LDN are associated with subsequent clinical response to anti-PD-1 therapy. To further explore this association, longitudinal analyses were performed in representative patients. Responders exhibited a lower or progressive decline in the frequency of CCR4^+^LDN during treatment that paralleled the sustained clinical benefit, whereas non-responders showed persistently elevated or increasing levels of these neutrophils, corresponding to disease progression and/or treatment failure (Figure 8D).

Although derived from a relatively small exploratory cohort that requires further validation in larger prospective studies, these data provide initial evidence that longitudinal monitoring of circulating CCR4^+^LDN may represent a useful blood-based biomarker for assessing early on-treatment response to anti-PD-1 therapy, with potential to guide treatment decisions and inform patient management in TNBC.

## Discussion

In this study, we establish the chemokine receptor CCR4 as a marker of a previously unrecognized subset of circulating LDN in BC. We demonstrate that CCR4^+^LDN expand during disease progression, display enhanced immunosuppressive and migratory properties, and are associated with adverse clinical outcomes and reduced response to PD-1 blockade. Collectively, these findings define a neutrophil population with direct clinical relevance and position CCR4 as a marker that refines the heterogeneous LDN compartment, supporting its potential as a biomarker and therapeutic target in BC.

Neutrophil heterogeneity has emerged as a major challenge in understanding the role of these cells in cancer (28,29). Although circulating LDN have consistently been associated with disease progression and poor prognosis in BC and other malignancies (18,19,30,31,32), their clinical translation has been limited by the lack of markers to distinguish detrimental neutrophils from the broader circulating neutrophil compartment (16). The identification of CCR4 in these cells represents a step towards addressing this limitation, defining a subset with distinct biological and clinical properties. To our knowledge, CCR4 expression has not previously been explored in neutrophils.

Notably, CCR4 is best known for mediating the recruitment of regulatory T cells through its ligands CCL17 and CCL22, contributing to immune suppression within the tumor microenvironment (33,34). Both increased Treg infiltration and elevated levels of CCL17 and CCL22 have been associated with poor prognosis in breast cancer (35,36). The identification of the same receptor in neutrophils suggests a convergence of adaptive and innate immune suppressive programs and raises the possibility that multiple immunosuppressive populations may exploit shared trafficking mechanisms to accumulate within the tumor microenvironment, thereby favoring immune escape.

Several observations from this study support the clinical relevance of this newly identified population. CCR4^+^LDN frequencies progressively accumulated with disease advancement, and elevated levels identified metastatic patients with shorter survival and more rapid disease progression.

Moreover, longitudinal analyses revealed that fluctuations in circulating CCR4^+^LDN closely paralleled changes in disease status, frequently accompanying or preceding clinical deterioration. Importantly, it is the rising frequency itself, rather than the baseline value, that carried the greatest prognostic weight.

Additionally, because LDN may accumulate in a variety of inflammatory conditions unrelated to cancer, CCR4 expression may provide greater specificity for identifying tumor-associated pathogenic neutrophils than total LDN frequencies alone. Supporting this interpretation, systemic concentrations of the CCR4 ligands CCL17 and CCL22, which are known to be secreted by tumor cells (37), were elevated in metastatic patients, consistent with the engagement of the CCR4-CCL17/CCL22 axis during BC progression.

Our findings further suggest that CCR4 expression may be regulated, at least in part, by tumor-associated signals. TGF-β is a well-established regulator of neutrophil polarization toward protumor phenotypes within the TME (27). Consistent with this role, metastatic patients exhibited elevated circulating TGF-β levels, which positively correlated with the frequency of CCR4^+^LDN. Moreover, exposure of HDN from BC patients to recombinant TGF-β increased CCR4 expression, whereas LDN remained unresponsive. This pattern aligns with previous reports showing that TGF-β promotes the conversion of HDN into LDN in tumor-bearing mice, suggesting that TGF-β may induce CCR4 expression in still-mature neutrophils (27), with full acquisition of the protumor, low-density phenotype requiring additional conditioning by the tumor microenvironment (38). Interestingly, TGF-β regulates CCR4 expression in Tregs through a FOXP3-dependent transcriptional program (26). Consistent with a similar mechanism in neutrophils, TGF-β stimulation in HDN was accompanied by increased FOXP3 and CCR4 expression, raising the possibility that neutrophils may exploit a comparable regulatory pathway during CCR4 acquisition. Further mechanistic studies will be required to define the molecular pathways regulating CCR4 expression in neutrophils, including the potential role of FOXP3 and the contribution of additional tumor-derived mediators, such as IL-10 and VEGF, alongside TGF-β, as described in Tregs (26).

Beyond their clinical associations, CCR4^+^LDN displayed a phenotypic profile consistent with a highly specialized protumor neutrophil population. Compared with CCR4^-^LDN, these cells exhibited significantly higher frequencies of cells positive for CD33, PD-L1, TLR4, CD36, LOX-1, CD98, and LAMP-1, together with higher arginase activity, indicating the simultaneous enrichment of multiple pathways associated with immune suppression, metabolic adaptation, cellular activation, and migration.

Elevated PD-L1 expression and arginase activity are consistent with their enhanced capacity to suppress T cell responses (39,40,41), whereas increased CD33 supports a less differentiated phenotype, in agreement with previous reports linking immature neutrophils to tumor-promoting functions (42). The enrichment of CD36, LOX-1, and CD98 further suggests that CCR4^+^LDN undergo metabolic reprogramming that may facilitate their persistence and function within the nutrient-deprived TME. CD36 has been implicated in lipid uptake and metabolic adaptation of immunosuppressive neutrophils (43), whereas LOX-1 is increasingly recognized as a hallmark of pathological neutrophils and polymorphonuclear myeloid-derived suppressor cells, promoting neutrophil survival, oxidative stress responses, angiogenesis, and metastatic dissemination (44). CD98 may similarly contribute to the bioenergetic capacity of CCR4^+^LDN. In systemic lupus erythematosus, CD98 on LDN mediates uptake of essential amino acids that fuel mitochondrial ATP production, particularly under glucose-limited conditions (45). In that context, CD98^+^LDN also produce more proinflammatory cytokines and chemokines than normal-density neutrophils and display resistance to apoptosis, contributing to tissue inflammation (45). A comparable role for CD98 in sustaining the metabolic and functional persistence of CCR4^+^LDN within the nutrient-deprived tumor microenvironment is therefore plausible and warrants further investigation. TLR4 signaling has been implicated in BC progression by promoting neutrophil activation, survival, and the production of tumor-supportive mediators, including APRIL, which enhances breast cancer cell proliferation and survival (46,47,48). Although APRIL was not evaluated in the present study, elevated TLR4 expression suggests that CCR4^+^LDN may be particularly responsive to tumor-derived danger signals, reinforcing their persistence and protumor functions. LAMP-1, together with CD98, has been implicated in integrin-mediated adhesion, cellular trafficking, and migratory responses (49,50,51). Consistent with their phenotypic profile, CCR4^+^LDN displayed enhanced CCR4-dependent migration toward CCL17, CCL22, and breast cancer cells, and were also detected within patient tumors. These observations support a model in which the CCR4-CCL17/CCL22 axis promotes the recruitment of this subset into the TME, thereby contributing to immune evasion and tumor progression.

This study also identifies CCR4^+^LDN as potent suppressors of antitumor immunity that may contribute to resistance to PD-1 blockade. Compared with CCR4^-^LDN, this subset more effectively inhibited T cell activation and impaired T cell-mediated tumor cell killing despite anti-PD-1 treatment, indicating that its immunosuppressive activity cannot be fully overcome by checkpoint inhibition alone. These findings align with accumulating evidence identifying neutrophils as key determinants of resistance to immune checkpoint inhibitors (52). Elevated circulating neutrophil counts and LDN frequencies have been associated with poor responses to immunotherapy across multiple tumor types (53,54,55). However, the specific neutrophil populations responsible for these effects have remained poorly defined. Our data extend this concept by identifying CCR4^+^LDN as a discrete immunosuppressive neutrophil subset that engages with multiple suppressive mechanisms beyond the PD-1/PD-L1 axis, likely including arginase-dependent pathways, metabolic adaptations, and additional immunoregulatory mediators, thereby limiting the efficacy of immune checkpoint blockade. The clinical relevance of these observations is further supported by our longitudinal pilot study of TNBC patients receiving anti-PD-1 therapy, in which early-on-treatment increases in CCR4^+^LDN frequencies were associated with poor clinical responses, whereas declining frequencies generally accompanied therapeutic benefit. Although these observations were obtained in a small exploratory cohort and require validation in larger prospective studies, they support the potential of CCR4^+^LDN as a dynamic biomarker for monitoring response to immune checkpoint blockade in BC.

The identification of CCR4^+^LDN also has important therapeutic implications. Unlike approaches aimed at broadly depleting neutrophils, which may compromise essential antimicrobial functions, targeting CCR4^+^ neutrophils offers the opportunity to selectively interfere with a pathological neutrophil subset while potentially preserving physiological neutrophil immunity. This concept is particularly attractive given the availability of clinically approved anti-CCR4 antibodies, such as mogamulizumab (56,57,58). Although these therapies were originally developed to target CCR4-expressing malignant cells and Tregs (59), our findings raise the possibility that CCR4^+^ neutrophils may represent an additional therapeutic target. Beyond repurposing existing agents, these findings may also support the development of novel anti-CCR4 antibodies or antibody fragments specifically optimized to target this neutrophil subset. Combining CCR4-directed therapies with immune checkpoint blockade could therefore simultaneously limit the recruitment of immunosuppressive neutrophils to the TME, while enhancing antitumor T cell responses, ultimately improving the efficacy of immunotherapy approaches.

Still, this study presents some limitations. The molecular mechanisms regulating CCR4 expression in neutrophils remain incompletely understood, and the immunotherapy cohort analyzed here was relatively small and exploratory. In addition, although our functional assays provide mechanistic insight into the immunosuppressive activity of CCR4^+^LDN, they do not fully recapitulate the complexity of the TME, and *in vivo* validation of CCR4 as a therapeutic target will be required to confirm its translational potential. Future studies combining single-cell transcriptomics, metabolic profiling, and more physiologically relevant experimental models, such as tumor-on-chip models and mouse models, will be important to define the developmental origin, molecular programs, and therapeutic vulnerabilities of this neutrophil subset.

In summary, we identify CCR4 as a marker of a previously unrecognized subset of immunosuppressive low-density neutrophils that expands during BC progression and displays enhanced migratory capacity, metabolic adaptation, and immunosuppressive activity associated with therapy resistance. By refining the heterogeneous LDN compartment into a functionally distinct population, CCR4^+^LDN offer new insight into neutrophil heterogeneity in cancer and represent a promising biomarker and therapeutic target for improving patient stratification, immunotherapy efficacy, and long-term outcomes in BC.

## Methods

### Sex as a biological variable

This study enrolled predominantly female patients, consistent with the well-established female predominance in BC epidemiology. A single male participant was included, however, given this limited representation, sex-stratified analyses were not performed. Therefore, whether these findings apply to male breast cancer remains to be determined.

### Study design and patient cohort

Peripheral blood from 198 BC patients (99 non-metastatic and 99 metastatic patients) was collected during routine clinical appointments at Instituto Português de Oncologia de Lisboa Francisco Gentil (IPO-Lisboa) and Hospital Professor Doutor Fernando Fonseca (HFF, Amadora, Portugal) (see cohort flowchart in Supplemental Figure 5). Whenever possible, follow-up samples were also collected, approximately every 3 months, with 46 metastatic BC patients included in the longitudinal study. Additionally, matched peripheral blood samples and surgical tumor specimens were obtained from 10 BC patients at HFF and Hospital CUF Tejo. Clinical and demographic characteristics of the study population are summarized in Table 1. Eligible participants were adults (≥ 18 years) with BC of any subtype who provided written informed consent before enrollment. Exclusion criteria included inability to provide informed consent, cognitive impairment, autoimmune disease, and pregnancy. Blood from 7 healthy individuals served as controls. Sample collection was performed during routine clinical visits and did not influence clinical management or treatment decisions.

**Table 1.**
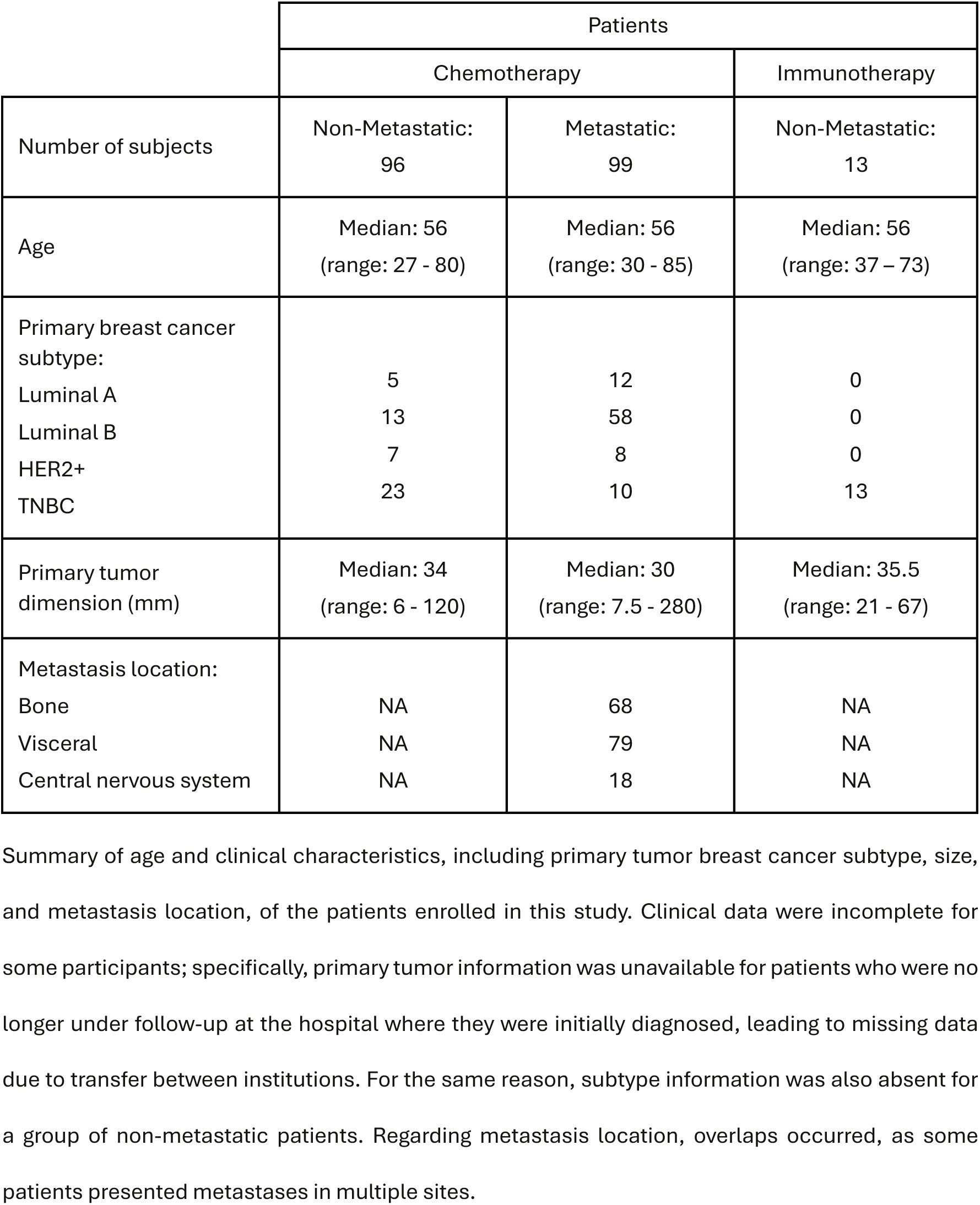
Characteristics of the breast cancer patients who contributed biological samples for this study.

### Processing of patient samples for experiments

Peripheral blood was collected into EDTA-containing Vacutainer tubes (BD Biosciences) and processed within 24 hours of collection. LDN, HDN, PBMCs, and plasma were isolated by density gradient centrifugation as previously described by our group^19^. Briefly, whole blood was layered over a Histopaque-1077/Histopaque-1119 bilayer (Sigma-Aldrich) and centrifuged without brake, allowing the separation of PBMCs, which may include circulating LDN, from the granulocyte fraction enriched in HDN. Both cellular fractions were used for immunophenotyping by flow cytometry and functional assays. Plasma was stored at -80°C for subsequent cytokine quantification by ELISA.

Tumor samples were mechanically dissociated using Medicons (BD Biosciences) to obtain single-cell suspensions. The resulting suspensions were filtered through a 30-μm mesh filter (Sysmex), washed with PBS, and prepared for flow cytometry analyses.

### Flow cytometry

Whole blood, isolated LDN and HDN fractions, and tumor-derived single-cell suspensions were stained with fluorochrome-conjugated monoclonal antibodies against surface and intracellular markers relevant to neutrophil and T cell phenotyping. For whole blood and HDN samples, erythrocytes were lysed with RBC Lysis Buffer (BioLegend). Intracellular staining was performed following fixation and permeabilization using the Fix/Perm kit (Invitrogen). Data acquisition, including CCR4 expression and the initial characterization of CCR4^+^ and CCR4^-^LDN, was performed on a BD FACSCanto II using FACSDiva software v8.0.1 (BD Biosciences). Following expansion of the antibody panel, additional phenotyping analyses were acquired on a BD FACSDiscover A8 using BD FACSChorus software v6.3 (BD Biosciences). All data were analyzed with FlowJo v10 (BD Biosciences).

LDN and HDN were identified as CD15^+^CD66b^+^ cells within the PBMC and granulocyte fractions, respectively. Tumor-associated neutrophils were identified as CD15⁺CD66b⁺ cells within tumor-derived single-cell suspensions. Cytotoxic T lymphocytes were identified as CD3^+^CD8^+^, helper T lymphocytes as CD3^+^CD4^+^, and regulatory T cells as CD4^+^CD25^high^CD127^low^ cells. The frequency of marker-positive cells was determined using gates established with unstained controls as references. A complete list of antibodies, fluorochromes, clones, and manufacturers is provided in Supplemental Table 4.

For experiments requiring isolated LDN and HDN populations, CD15^+^CD66b^+^ neutrophils were sorted by fluorescence-activated cell sorting (FACS; BD FACSAria III) from the PBMC and/or granulocyte fractions, respectively, obtained from BC patients. For co-culture experiments, the corresponding CD15-depleted PBMC fraction, containing lymphocytes and monocytes, was collected in parallel.

### Cell culture

The human TNBC cell line MDA-MB-231 was cultured in Dulbecco’s Modified Eagle Medium (DMEM; Gibco) supplemented with 10% FBS (Biowest) and 1% Penicillin/Streptomycin (GE Healthcare) at 37°C in a humidified 5% CO₂ atmosphere. Cells were routinely passaged at 80–90% confluency and used for the 2D and 3D co-culture experiments with patient-derived immune cells. Cells tested negative for mycoplasma contamination before use in the experiments.

### ELISA

Plasma concentrations of CCL17, CCL22, and TGF-β were measured in BC patients and healthy donors using commercially available ELISA kits, following the manufacturers’ instructions. Details of the kits used are provided in Supplemental Table 5. Absorbances at 450 nm with wavelength correction at 570 nm using a Synergy HT Multi-Detection Microplate Reader (BioTek), and analyte concentrations were interpolated from a 4-parameter logistic standard curve.

### Arginase activity

Arginase activity was determined in supernatants from CCR4^+^ and CCR4^-^LDN cultures using the Arginase Activity Assay Kit (Sigma-Aldrich, Supplemental Table 6) according to the manufacturer’s instructions. Absorbance was measured at 430 nm using a Synergy HT Multi-Detection Microplate Reader (BioTek), and results were expressed as nmol urea/minute/mL.

### TGF-β effects on CCR4 expression and FOXP3 localization

PBMC and granulocyte fractions isolated from metastatic BC patients and healthy donors were cultured overnight at 37°C and 5% CO₂ with recombinant human TGF-β1 (1 or 10 ng/mL; PeproTech, Supplemental Table 6) or vehicle control. Cells were subsequently stained with anti-CD15 and anti-CCR4 antibodies, and CCR4 expression was assessed by flow cytometry. CCR4 expression was quantified as the ratio of CCR4 mean fluorescence intensity (MFI) in CD15^+^CCR4^+^ cells relative to CD15^+^CCR4^-^cells, which served as an internal normalization control.

To further investigate the relationship between CCR4 and FOXP3 expression at the single-cell level, neutrophils were stained with anti-CD66b, anti-CCR4, and anti-FOXP3 antibodies and analyzed using a BD FACSDiscover™ A8 Cell Analyzer. High-content cellular images were acquired simultaneously with flow cytometry and analyzed using FlowJo v10 to evaluate the co-expression and intracellular distribution of CCR4 and FOXP3. Details of all antibodies are provided in Supplemental Table 4.

### Transwell migration assays

Neutrophil migration was assessed using 5-μm pore transwell inserts (Sarstedt) to evaluate CCR4-dependent chemotaxis toward chemokine and tumor-derived stimuli, using sorted neutrophils from patient blood.

For chemokine-driven migration assays, recombinant human CCL17 (10 ng/mL) or CCL22 (100 ng/mL) (PeproTech, Supplemental Table 6) was added to the lower chamber, while medium alone was used as a control. Migrated neutrophils were collected and quantified by flow cytometry using anti-CD15 and anti-CCR4 antibodies and counting beads. Results were expressed as the percentage of migrated CD15^+^CCR4^+^ cells relative to the control condition.

For cancer cell-directed migration assays, MDA-MB-231 cells were seeded in the lower chamber 24h before the assay. Sorted CD15^+^CD66b^+^ LDN were added to the upper chamber, and migrated cells were quantified by flow cytometry. Results were expressed as the fold change in migration toward tumor cells relative to the controls.

Both experiments were repeated with neutrophils pre-incubated with either a CCR4-blocking antibody (10 μg/mL) or an isotype control antibody before migration.

### LDN-T cell co-culture and checkpoint blockade

CCR4^+^CD15^+^LDN, CCR4^-^CD15^+^LDN, and remaining PBMCs (CD15^-^) were sorted from PBMCs of metastatic BC. CD3^+^ T cells were cultured alone or co-cultured with CCR4^+^ or CCR4^-^LDN at a 1:1 ratio and stimulated with PMA (35 ng/mL) and ionomycin (1 µg/mL). T cell activation was assessed by flow cytometry analysis of CD25 and CD69 expression within CD3^+^ T cells. For experiments with immune checkpoint blockade, anti-PD-1 antibody (10 μg/mL) was added to the LDN–T cell co-cultures prior to stimulation, and T cell activation was evaluated after 24h by flow cytometry.

### Tumor spheroid cytotoxicity assay

MDA-MB-231 tumor spheroids were generated by adapting a protocol previously established by our group (60) and co-cultured with activated T cells previously exposed to CCR4^+^ or CCR4^-^LDN. Tumor cell viability was assessed as a functional readout of T cell–mediated cytotoxicity. For checkpoint blockade experiments, anti-PD-1 antibody was added to spheroid co-cultures, and MDA-MB-231 viability was assessed after 24h using a Fixable Viability Dye (BD Biosciences) and analyzed by flow cytometry.

## Statistics

Statistical analysis was performed in GraphPad Prism v10.3.1, and results were considered statistically significant for p-values < 0.05. Depending on the experimental design and data distribution, comparisons between groups were conducted using non-parametric tests (Mann-Whitney, Kruskal-Wallis, Wilcoxon, Friedman) or parametric tests (Welch, one-way ANOVA, paired t-test), as appropriate. Categorical variables were analyzed with the Chi-squared test. Correlations were assessed using Spearman’s rank correlation. A receiver operating characteristic (ROC) curve analysis using survival data from metastatic BC patients was performed to define the optimal cut-off value for CCR4^+^LDN frequencies, maximizing sensitivity to minimize false negatives. The resulting threshold was applied to analyses of disease progression, survival, and swimmer plots. Survival distributions were compared using the Gehan-Breslow-Wilcoxon test. Data are presented as mean ± SD. All experiments included at least three independent biological replicates. For *in vitro* assays, neutrophils and PBMCs were obtained from different donors, and each biological replicate corresponded to an independent donor sample to account for inter-donor variability.

## Study approval

This study was approved by the Ethics Committees of Hospital Prof. Doutor Fernando da Fonseca, Instituto Português de Oncologia de Lisboa Francisco Gentil, Hospital de Santa Maria, Hospital de Vila Franca de Xira, Instituto CUF Oncologia, and NOVA Medical School. All procedures involving human participants were performed in accordance with the ethical standards of the participating institutions and with the principles of the Declaration of Helsinki. Written informed consent was obtained from all participants at the time of enrollment and before the collection of biological samples and clinical data. The study complied with applicable national legislation governing the use of human biological samples. Clinical data were pseudoanonymized and managed in accordance with the General Data Protection Regulation (EU), and all study records were securely maintained.

## Data availability

The data supporting the findings of this study are available in the article, its Supplemental Material, and in the Supporting Data Values file, in accordance with the journal’s data availability policy. Additional information required to reproduce or reanalyze the results reported in this paper is available from the corresponding author upon reasonable request.

## Author contributions

BFC and DG contributed equally to this work. Both conducted all the experiments, analyzed and interpreted the data, performed the statistical analysis, assembled all the figures, and wrote the manuscript. RS, IB, AL, MP, BM, DPS, and NDS helped in the execution of flow cytometry and cell culture experiments. SB, TM, MV, GV, MM, MGG, and SCV were responsible for selecting the patients, collecting the samples and clinical data, and contributing to the scientific discussion. JMM contributed to the statistical analysis. AJ contributed to the scientific discussion. MGC designed and supervised the study, contributed to data interpretation, and participated in the writing and revision of the manuscript. All the authors revised and approved the final manuscript.

## AI disclosure

AI tools were used to assist with revision of selected sections of the manuscript and to support the preparation and refinement of illustrative elements included in the figures. The authors ensured the integrity and accuracy of AI output presented in this article.

## Authorship notes

BFC and DG are co-first authors.

## Acknowledgements

We sincerely thank all breast cancer patients who generously participated in this study. We are deeply grateful to the nurses, oncologists, and clinical teams from the participating hospitals for their invaluable collaboration in patient recruitment and in the collection of blood samples, tumor specimens, and clinical data. We also acknowledge the support of the Flow Cytometry and Cell Culture Facilities of NOVA Medical School, whose technical expertise was essential to this work. Finally, we thank the Research Unit iNOVA4Health and the Associated Laboratory LS4FUTURE for providing the scientific environment and infrastructure that supported this research.

