## Supplemental Material for "CCR4^+^low-density neutrophils define a novel immunosuppressive subset associated with breast cancer progression and early mortality"

### Supplemental Figures:

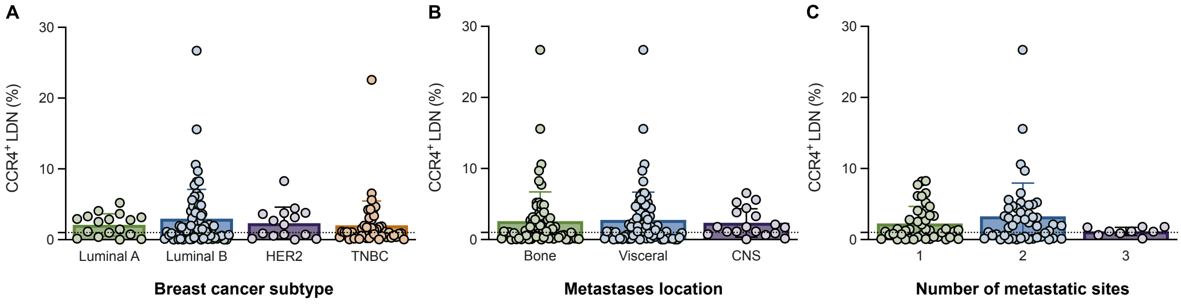


**Supplemental Figure 1 - The frequency of CCR4⁺ low-density neutrophils in breast cancer patients varies across all subtypes, metastatic sites, and number of metastases. (A)** Percentage of CCR4⁺ low-density neutrophils (CCR4⁺LDN) in the blood of patients with non-metastatic and metastatic breast cancer (BC), categorized by subtype. Luminal A (green, *n* = 17), Luminal B (blue, *n* = 71), HER2 (purple, *n* = 15), or Triple-Negative (TNBC, orange, *n* = 46). **(B)** Percentages of CCR4⁺LDN present in the blood of BC patients presenting with bone (green, *n* = 68), lung (blue, *n* = 79), central nervous system (CNS, purple, *n* = 18). **(C)** Percentages of CCR4⁺LDN present in the blood of BC patients presenting with 1 (green, *n* = 44), 2 (blue, *n* = 47), or 3 (purple, *n* = 9) metastatic sites.

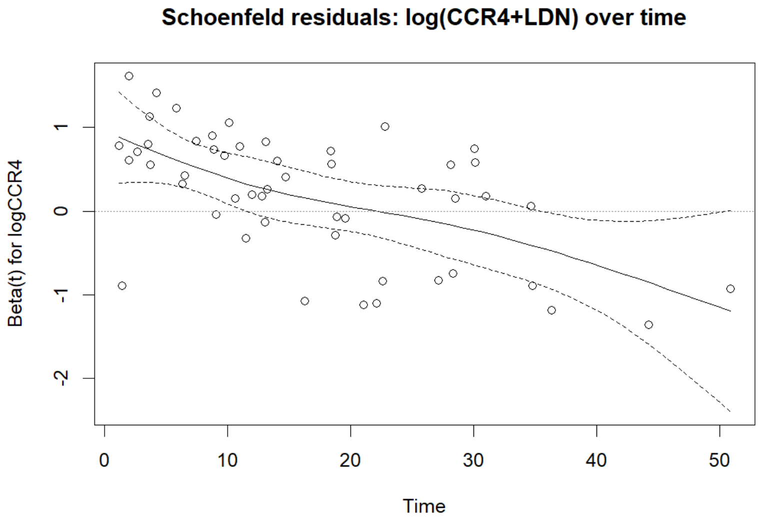


**Supplemental Figure 2 - Scaled Schoenfeld residuals plot for log(CCR4$^+$LDN) over time.** The solid line represents the smoothed spline fit, and the dashed lines represent the 95% confidence intervals. The downward trend indicated a time-dependent effect for this covariate, with the proportional-hazards test rejecting constancy ($p$ = 0.0007).

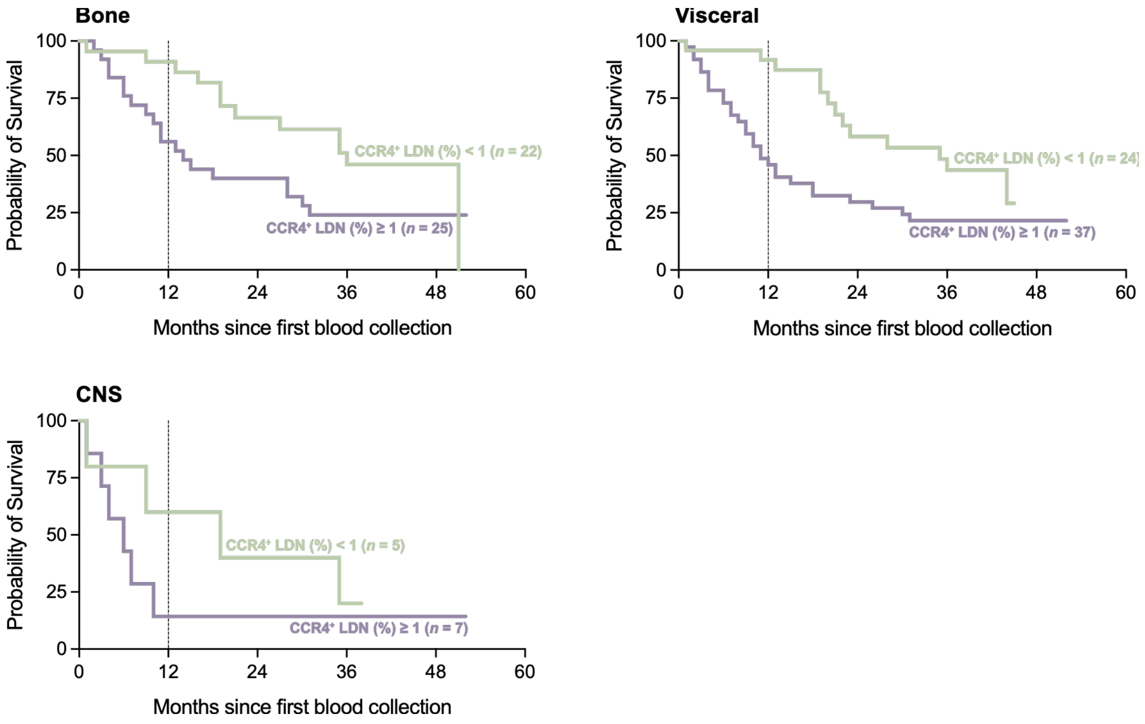


**Supplemental Figure 3 - Increased frequencies of CCR4$^+$ low-density neutrophils are associated with a worse prognosis, reflected in reduced survival, even after stratification by metastatic site.** Probability of survival of mBC patients exhibiting less than 1% (green) and patients exhibiting 1% or more (purple) of CCR4$^+$LDN in their blood, stratified by metastatic site: bone metastasis (HR: 0.4899 (95% CI, 0.24-0.99), *$p$ = 0.0108); visceral metastasis (including lung, liver, ganglionar, and others) (HR: 0.4442 (95% CI, 0.24-0.81), **$p$ = 0.0012); and central nervous system (CNS) metastasis (HR: 0.5605 (95% CI, 0.16-1.95), $p$ = 0.2958). Statistical analysis was performed with the Gehan-Breslow-Wilcoxon test.

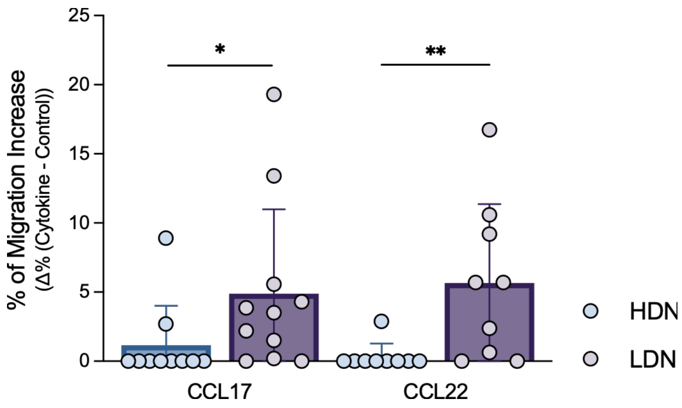


**Supplemental Figure 4 – Compared to high-density neutrophils, low-density neutrophils exhibited enhanced migratory ability towards CCL17 and CCL22.** Percentage increase in migration of high-density neutrophils (HDN, blue, $n$ = 10) and low-density neutrophils (LDN, purple, $n$ = 11) towards CCL17 and CCL22, normalized to the control. Statistical analysis was determined with the Mann-Whitney test. *$p < 0.05$, ** $p < 0.01$.

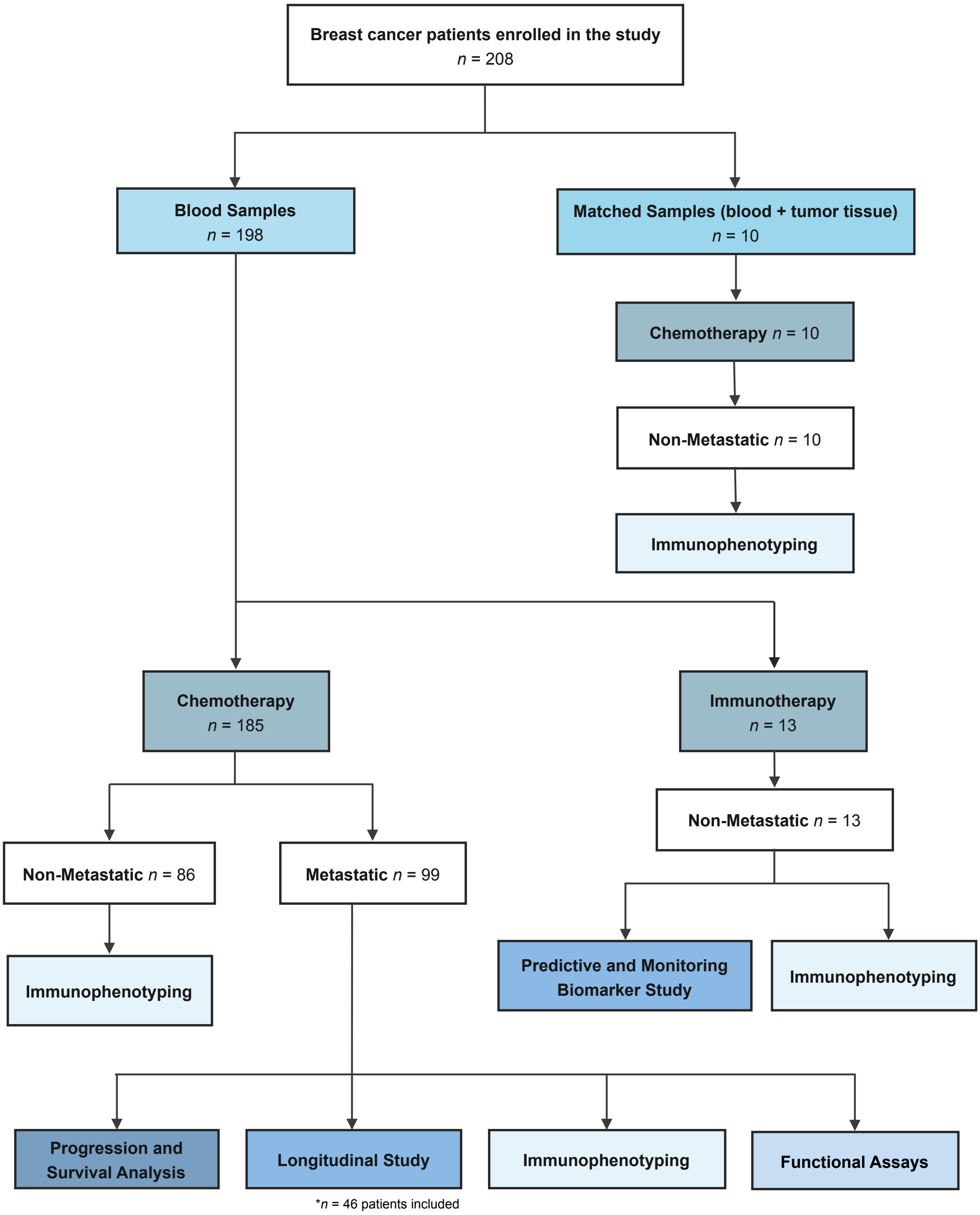


**Supplemental Figure 5 – Flowchart of the breast cancer patients enrolled in this study.** Blood samples from 208 breast cancer (BC) patients were analyzed, including 198 blood and 10 matched (blood + tumor tissue) samples, from non-metastatic or metastatic patients undergoing chemotherapy or immunotherapy. Neutrophil frequency and their respective immunophenotypes were assessed in both cohorts. Neutrophils derived from metastatic patients were further used for progression and survival analyses and for functional assays. For the longitudinal study, 46 of the 99 metastatic patients were included. The remaining 53 patients were excluded due to insufficient clinical data, widely spaced follow-up samples, or hospital transfer.

### Supplemental Tables:

**Supplemental Table 1 - Primary multivariable Cox model (*n* = 75, deaths = 49).** Each HR is adjusted for all the others.

| Term | coef (log-HR) | HR | 95% CI | *p* |
|---|---|---|---|---|
| CCR4$^+$LDN (*per* log unit) | 0.212 | 1.24 | 0.98–1.56 | 0.073 |
| Lead-time, months (metastasis → draw) | 0.008 | 1.01 | 1–1.02 | 0.095 |
| Age (*per* year) | 0.017 | 1.02 | 0.99–1.05 | 0.253 |
| Subtype: HER2+ (*vs* HR+/HER2-) | -0.537 | 0.58 | 0.27–1.26 | 0.172 |
| Subtype: TNBC (*vs* HR+/HER2-) | 1.475 | 4.37 | 1.34–14.2 | 0.014 |
| Visceral involvement (yes *vs* no) | 0.042 | 1.04 | 0.46–2.38 | 0.921 |
| Number of metastatic sites | 0.398 | 1.49 | 0.85–2.61 | 0.164 |
| Relapsed (*vs* de-novo metastatic) | -0.106 | 0.90 | 0.46–1.75 | 0.754 |
| LDH (*per* log unit) | 0.509 | 1.66 | 1.1–2.51 | 0.015 |

**Supplemental Table 2 - Primary multivariable Cox model (*n* = 75, deaths = 49).** Each HR is adjusted for all the others.

| Term | coef (log-HR) | HR | 95% CI | *p* |
|---|---|---|---|---|
| CCR4$^+$LDN per log unit — ≤ 12 months | 0.795 | 2.21 | 1.45–3.38 | <0.001 |
| CCR4$^+$LDN per log unit — >12 months | -0.089 | 0.91 | 0.71–1.18 | 0.487 |
| CCR4$^+$LDN ≥ 1% — ≤ 12 months | 1.626 | 5.08 | 1.5–17.2 | 0.009 |
| CCR4$^+$LDN ≥ 1% — >12 months | -0.142 | 0.87 | 0.42–1.81 | 0.705 |

**Supplemental Table 3 – Comparison of time-varying vs baseline-only Cox models (49 deaths).** Results demonstrate that the current (updated) value of the covariate significantly predicts mortality risk, whereas the enrolment (baseline) value alone does not.

| Model | HR | 95% CI | *p* | C-index |
|---|---|---|---|---|
| Time-varying: current log(CCR4$^+$LDN) | 1.60 | 1.28–2.01 | <0.001 | 0.707 |
| Time-varying: current CCR4$^+$LDN ≥1% | 4.49 | 2.17–9.28 | <0.001 | 0.707 |
| Baseline only: enrolment log(CCR4$^+$LDN) | 1.19 | 0.97–1.46 | 0.092 | 0.632 |

**Supplemental Table 4 – Antibodies and dyes used for flow cytometry.**

| Antibodies/Dyes | Clone | Manufactor | Cat # |
|---|---|---|---|
| anti-CD3-APC | UCHT1 | BioLegend | 300412 |
| anti-CD4-FITC | OKT4 | BioLegend | 317408 |
| anti-CD8-PacificBlue | HIT8a | BioLegend | 300928 |
| anti-CD8-PE | HIT8a | BioLegend | 300908 |
| anti-CD11b-FITC | ICRF44 | BioLegend | 301330 |
| anti-CD15-PE | HI98 | BioLegend | 301906 |
| anti-CD25-PE | BC96 | BioLegend | 302606 |
| anti-CD33-APC-Cy7 | P67.6 | BioLegend | 366614 |
| anti-CD36-APC-Cy7 | 5-271 | BioLegend | 336214 |
| anti-CD45-PerCP | HI30 | BioLegend | 304026 |
| anti-CD66b-APC | G10F5 | BioLegend | 305118 |
| anti-CD69-PerCP | FN50 | BioLegend | 310928 |
| anti-CD98-FITC | MEM-108 | BioLegend | 315603 |
| anti-CD127-PE/Cy7 | A019D5 | BioLegend | 986008 |
| anti-CCR4-BV421 | L291H4 | BioLegend | 359414 |
| anti-CCR4-RB780 | 1G1 | BD Biosciences | 755494 |
| anti-FOXP3-RB613 | 236A-E7 | BD Biosciences | 571236 |
| anti-HLA-DR-APC | L243 | BioLegend | 307610 |
| anti-LAMP-1-PE/Cy | H4A3 | BioLegend | 328608 |
| anti-LOX-1-APC | 15C4 | BioLegend | 358606 |
| anti-MD-1-FITC | F-5 | Santa Cruz Biotechnology | sc-390613 |
| anti-MMP-9-FITC | E-11 | Santa Cruz Biotechnology | sc-393859 |
| anti-MPO | 266-6K1 | Santa Cruz Biotechnology | sc-52707 |
| anti-N. Elastase-AlexaFluor 647 | G-2 | Santa Cruz Biotechnology | sc-55549 |
| anti-PD-L1-APC | 29E.2A3 | BioLegend | 329708 |
| anti-TLR4-PE | HTA125 | BioLegend | 312806 |
| anti-VEGF | C-1 | Santa Cruz Biotechnology | sc-7269 |
| Anti-mouse Alexa Fluor 488 | - | Invitrogen | A32723TR |
| Fixable Viability Stain 450 | - | BD Biosciences | 562247 |
| Zombie Aqua Fixable Viability Dye | - | BioLegend | 423101 |

**Supplemental Table 5 – ELISA and enzymatic kits.**

| Kits | Manufactor | Cat # |
|---|---|---|
| Human CCL17/TARC DuoSet ELISA | R&D System | DY364 |
| Human CCL22/MDC DuoSet ELISA | R&D System | DY336 |
| DuoSet ELISA Ancillary Reagent Kit 2 | R&D System | DY008B |
| LEGEND MAX™ Total TGF-β1 ELISA Kit | BioLegend | 436707 |
| Arginase Activity Kit | Sigma-Aldrich | MAK112 |

**Supplemental Table 6 – Recombinant proteins.**

| Recombinant Protein | Manufactor | Cat # |
|---|---|---|
| Human CCL17 (TARC) | PeproTech | 300-30-20UG |
| Human CCL22 (MDC) | PeproTech | 300-36A-20UG |
| Human TGF-β1 | PeproTech | 100-21-10UG |
